# Synthetic Cannabidiol Attenuates Heart Failure Progression with Concomitant and Post Injury Administration Through Modulation of Immune and Endothelial to Mesenchymal Transition – Related Remodeling Programs

**DOI:** 10.64898/2026.07.10.737555

**Authors:** Muthu Kumar Krishnamoorthi, Abraham Mendez-Fernandez, Khush Patel, Rajarajan Amirthalingam Thandavarayan, Gerardo Garcia Rivas, Omar Lozano Garcia, Kartiga Natarajan, Mahwash Kassi, Rayan Yousefzai, Guillermo Torre-Amione, Victor Manuel Trevino Alvarado, Arvind Bhimaraj

## Abstract

**Background:** Cardiac fibrosis is a central driver of adverse remodeling in heart failure with reduced ejection fraction (HFrEF), yet therapies directly targeting these pathways remain less established. We investigated the role of a pharmaceutical-grade synthetic (s) cannabidiol in HFrEF using an invitro and in vivo strategy.

**Methods:** HFrEF was induced in 12-week-old C57BL/6J mice using angiotensin II, L-NAME, and salt exposure. A 5-week (w) s-cannabidiol course was administered either concomitantly (beginning at week 0) during disease induction or after disease induction (beginning at week 4). Echocardiography, cardiac morphological characterization was performed at 5 and 9 weeks of the experiment. Cardiac tissue was processed for RNA extraction. Standard statistical and informatics methodology was used to compare groups.

**Results:** At 5 weeks, mice in the concomitant s-cannabidiol group had reduced cardiomyocyte hypertrophy and fibrosis area with better isovolumetric relaxation time, ejection fraction, and fractional shortening compared to HFrEF mice. In the treatment after disease induction model, at 9 weeks, s-cannabidiol treated mice maintained therapeutic effect compared to 5w HFrEF mice but also had enhanced structural and functional recovery compared to mice that recovered naturally. Bulk RNA-sequencing analysis demonstrated a significant transcriptional change in HFrEF compared to controls, with s-cannabidiol partially shifting the cardiac transcriptome away from the failing state and attenuates the HF-enriched transcriptional programs of oxidative stress, inflammatory signaling, hypoxia, apoptosis, p53/MYC/mTORC1/E2F remodeling, and EMT/fibrotic remodeling, while enriching lipid/peroxisomal metabolic pathways. In an invitro HUVEC model of Endothelial to Mesenchymal Transition (EndMT), s-cannabidiol inhibited the transition and also reversed established EndMT, with these effects attenuated by pharmacologic inhibition of CB2 and PPARγ, but not CB1 receptors.

**Conclusions:** s-cannabidiol attenuates adverse remodeling in experimental HFrEF, promotes recovery after injury, and is associated with suppression of EndMT-related programs mediated through CB2/PPARγ-linked endothelial signaling.

## INTRODUCTION

Heart failure (HF) remains a major contributor to cardiovascular morbidity and mortality worldwide.^1^ Therapeutics to treat Heart failure with reduced ejection fraction (HFrEF) have evolved through various prevailing pathobiological understandings of the disease state ranging from volume overload, neurohormonal activation, hemodynamic energetics, etc. Most of these have been cardiomyocyte centric but such a view has been challenged by recent revelations of a complex cellular milieu of the heart with the discoveries of an important role of non-cardiomyocytes.^2^ Interstitial fibrosis is mediated by non-cardiomyocytes and is a central determinant of adverse remodeling and disease progression in HF.^3^ Excessive extracellular matrix deposition disrupts myocardial architecture, impairs myocardial compliance, and contributes to progressive cardiac dysfunction due to maladaptive persistence. ^3^

Cannabidiol, a non-psychotropic phytocannabinoid, has shown anti-inflammatory and anti-fibrotic^4^ effects in murine models of heart failure has recently shown to attenuate fibrosis, limit pathological remodeling, and preserve cardiomyocyte health, specifically through a mitochondrial protection in cardiomyocytes.^5, 6^ However, the mechanisms underlying such anti-fibrotic impact on the heart remain poorly understood. In particular, whether cannabidiol acts only through direct effects on cardiomyocytes or also modulates non-myocyte cellular processes central to myocardial fibrosis^2, 7^ has not been established.

We therefore hypothesized that cannabidiol exerts both preventive and therapeutic effects in HF and that its anti-fibrotic actions are mediated through direct actions on non-cardiomyocyte population of cells. To test this hypothesis, we used a murine model of non-ischemic HFrEF and a murine model of natural recovery following HFrEF induction^8, 9^. Using pharmaceutical-grade synthetic cannabidiol (s-cannabidiol) administered subcutaneously, together with bulk RNA-sequencing of cardiac tissue and receptor-blocking experiments, we examined the effects of cannabidiol when given concomitantly during disease induction and later after induction of HFrEF injury. With a signal from the transcriptomics showing a role in Epithelial/Endothelial to mesenchymal transition (E/EndMT)-related programs, we then used an invitro model of EndMT^10^ to analyze the direct impact of cannabidiol on endothelial phenotype and candidate receptor pathways. Our findings show that s-cannabidiol has a protective effect in HFrEF with a multi-axis cardioprotective mechanism in which it reduces inflammatory-fibrotic injury and stress remodeling while partially restoring metabolic homeostasis. The antifibrotic mechanism is partly mediated through suppression of EndMT via cannabinoid receptor 2 (CB2) and peroxisome proliferator-activated receptor-γ (PPARγ) signaling on endothelial cells.

## METHODS

### Data Availability Statement

A detailed description of the methods and supporting data is available in the Supplementary Material. Supporting source data, processed data files, and analytic outputs used for the transcriptomic analyses are provided in the Supplementary Material and accompanying data files. Bulk RNA-sequencing data generated in this study can be made available on request at the discretion of the authors. Related materials, additional data, and analytic techniques are available from the corresponding author upon reasonable request.

### Animals and study protocol

Twelve-week-old male C57BL/6J mice (Inotiv, USA) were housed under controlled temperature with a 12-hour light/dark cycle and provided standard rodent chow and water ad libitum. After a 1-week acclimatization period, mice were randomly assigned to one of the experimental groups described below **(Supplementary Figure 1)**. HFrEF was induced by administration of L-NAME (0.3 mg/mL) and 1% NaCl in drinking water ad libitum, followed by implantation of a subcutaneous osmotic minipump (ALZET model 1004, DURECT Corporation, Cupertino, CA) delivering angiotensin II (0.7 mg/kg/day) at 1 week which lasts for 4 weeks. Mice were anesthetized with inhaled isoflurane during pump implantation. At the end of the *5-week induction phase*, mice developed a phenotype consistent with HFrEF, including cardiac hypertrophy, fibrosis, and reduced systolic function. For the post-induction recovery experiment, L-NAME treatment was discontinued at week 5, and mice were maintained until week 9 without exposure to L-NAME or angiotensin II. Mice were euthanized according to institutional protocol at 5 and 9 weeks in the respective experiments. Age-matched controls were used for all comparisons. Final comparisons were performed between the following: (a) Week 5 control (W5 Ctrl); (b) W5 Ctrl with **c**oncomitant **s**ynthetic cannabidiol (c-s-cannabidiol) injections (W5 Ctrl+c-s-cannabidiol); (c) W5 HFrEF; (d) W5 HFrEF +c-s-cannabidiol; (e) W9 Ctrl; (f) W9 mice after HFrEF (5 week induction) followed by 4 weeks of natural recovery (W9-HFrEF+Recovery) and (g) W9 mice after HFrEF induction followed by treatment of s-cannabidiol (from week 4 to week 9 of the experiment) administered post/induction (pi) (W9 HFrEF+pi-s-cannabidiol). Unless otherwise specified, each experimental group contained n≥5 mice **(Supplementary Figure 1)**.

### Statistical analysis

Other statistical analyses not involving RNA-Seq data were performed using GraphPad Prism version 10 (GraphPad Software, San Diego, CA). Normality was assessed using the Shapiro–Wilk test. For comparisons among more than two groups, normally distributed data were analyzed using 1-way analysis of variance (ANOVA) followed by Tukey’s multiple-comparisons test. When data were not normally distributed, comparisons were performed using the Kruskal–Wallis test followed by Dunn’s multiple-comparisons test. For two-group comparisons, normally distributed data were analyzed using an unpaired 2-tailed Student’s t test, whereas non-normally distributed data were analyzed using the Mann–Whitney U test. A 2-sided P<0.05 was considered statistically significant. Data are presented as mean ± SEM, unless otherwise specified.

## RESULTS

### S-cannabidiol attenuates cardiac structural and functional deterioration when administered concomitantly with HFrEF induction agents

To determine the impact of s-cannabidiol preventing the onset of HFrEF, we administered s-cannabidiol concomitantly with disease induction agents, comparing W5 control, W5 HFrEF, and W5 HFrEF+c-s-cannabidiol groups **(Supplementary Figure 1)**. W5 HFrEF mice developed marked adverse cardiac remodeling, evidenced by increased cardiomyocyte size and fibrosis compared to controls, with a significant attenuation of these changes in s-cannabidiol treated animals **(Figure 1)**. Functional assessment by echocardiography showed that HFrEF induction was associated with a prolonged isovolumetric relaxation time (IVRT), reduction in ejection fraction (EF) and fractional shortening (FS) consistent with impaired diastolic and systolic function while these abnormalities were improved by concomitant s-cannabidiol administration **(Figure 1G-I)**. Collectively, these data indicate that concomitant s-cannabidiol administration attenuates adverse hypertrophic and fibrotic remodeling while preserving both diastolic and systolic function during HFrEF development.

**Figure 1.**
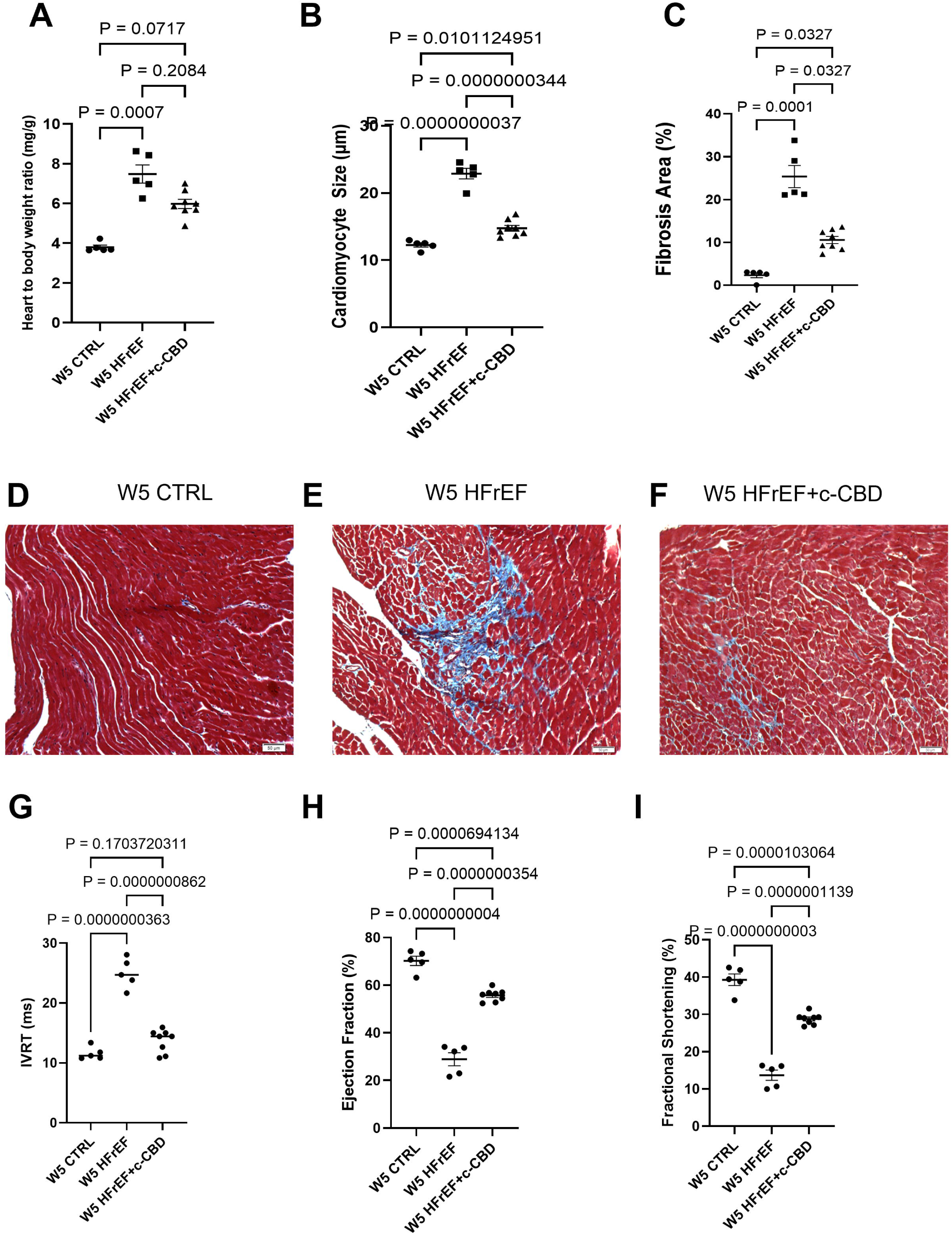
Concomitant synthetic cannabidiol treatment attenuates adverse cardiac remodeling and preserves ventricular function in experimental HFrEF. **A**, Heart weight-to-body weight ratio (HW/BW), demonstrating increased cardiac hypertrophy in W5 HFrEF mice and attenuation with concomitant synthetic cannabidiol treatment. **B**, Quantification of cardiomyocyte size, showing HFrEF-associated cardiomyocyte hypertrophy and its reduction with concomitant synthetic cannabidiol treatment. **C**, Quantification of myocardial fibrotic area from Masson trichrome-stained sections, demonstrating increased collagen deposition in HFrEF and reduced fibrosis with concomitant synthetic cannabidiol treatment. **D–F**, Representative Masson trichrome-stained myocardial sections from W5 CTRL (**D**), W5 HFrEF (**E**), and W5 HFrEF treated concomitantly with synthetic cannabidiol (**F**). Collagen is shown in blue and myocardium in red. Scale bar, 50 μm. **G**, Isovolumetric relaxation time, demonstrating impaired diastolic relaxation in HFrEF and improvement with concomitant synthetic cannabidiol treatment. **H**, Left ventricular ejection fraction (EF), showing systolic dysfunction in HFrEF and improvement with concomitant synthetic cannabidiol treatment. **I**, Fractional shortening (FS), demonstrating reduced systolic performance in HFrEF and improvement with concomitant synthetic cannabidiol treatment. For **A** and **C**, groups were compared using the Kruskal–Wallis test followed by Dunn multiple-comparisons test. For **B** and **G– I**, groups were compared using 1-way ANOVA followed by Tukey multiple-comparisons test. Data are presented as mean±SEM. *P*<0.05 was considered statistically significant. W5 CTRL indicates week 5 control; W5 HFrEF, week 5 heart failure with reduced ejection fraction; W5 HFrEF+c-CBD, week 5 heart failure with reduced ejection fraction treated with concomitant treatment of synthetic cannabidiol; and EF, ejection fraction

### S-cannabidiol enhances structural and functional recovery when administered after HFrEF induction

To evaluate whether s-cannabidiol retains therapeutic benefit after injury has begun, we used a post-induction (pi) recovery model in which HFrEF-inducing stimuli were withdrawn at week 5, followed by either passive observation of mice or pi-s-cannabidiol treatment **(Supplementary Figure 1)**. Relative to the established W5 HFrEF state, W9 HFrEF+pi-s-cannabidiol mice showed lower heart weight-to-body weight ratio, smaller cardiomyocyte size, reduced fibrotic burden, and improved IVRT, ejection fraction, and fractional shortening, indicating that delayed cannabidiol treatment ameliorated key structural and functional features of HFrEF **(Supplementary Figure 2)**.

Mice that were observed passively (natural recovery) after the HFrEF induction for 5 weeks did show improvement of their cardiac HF parameters at Week 9 (data not shown). The W9 HFrEF+Recovery and W9 HFrEF+pi-s-cannabidiol groups were compared to determine whether active treatment conferred benefit beyond passive recovery. Mice treated with s-cannabidiol exhibited better structural remodeling than animals undergoing natural recovery alone. The heart-to-body weight ratio, cardiomyocyte size and fibrotic area were significantly lower in the s-cannabidiol-treated group (W9 HFrEF+pi-s-cannabidiol). **(Figure 2 A-D)** These structural improvements were accompanied by a significantly better diastolic performance reflected by a shorter IVRT **(Figure 2E**). The changes in EF and FS were not significant **(Figure 2F & G)**. Of note, systolic indices in the week-9 natural recovery group had substantially normalized by this time point.

**Figure 2.**
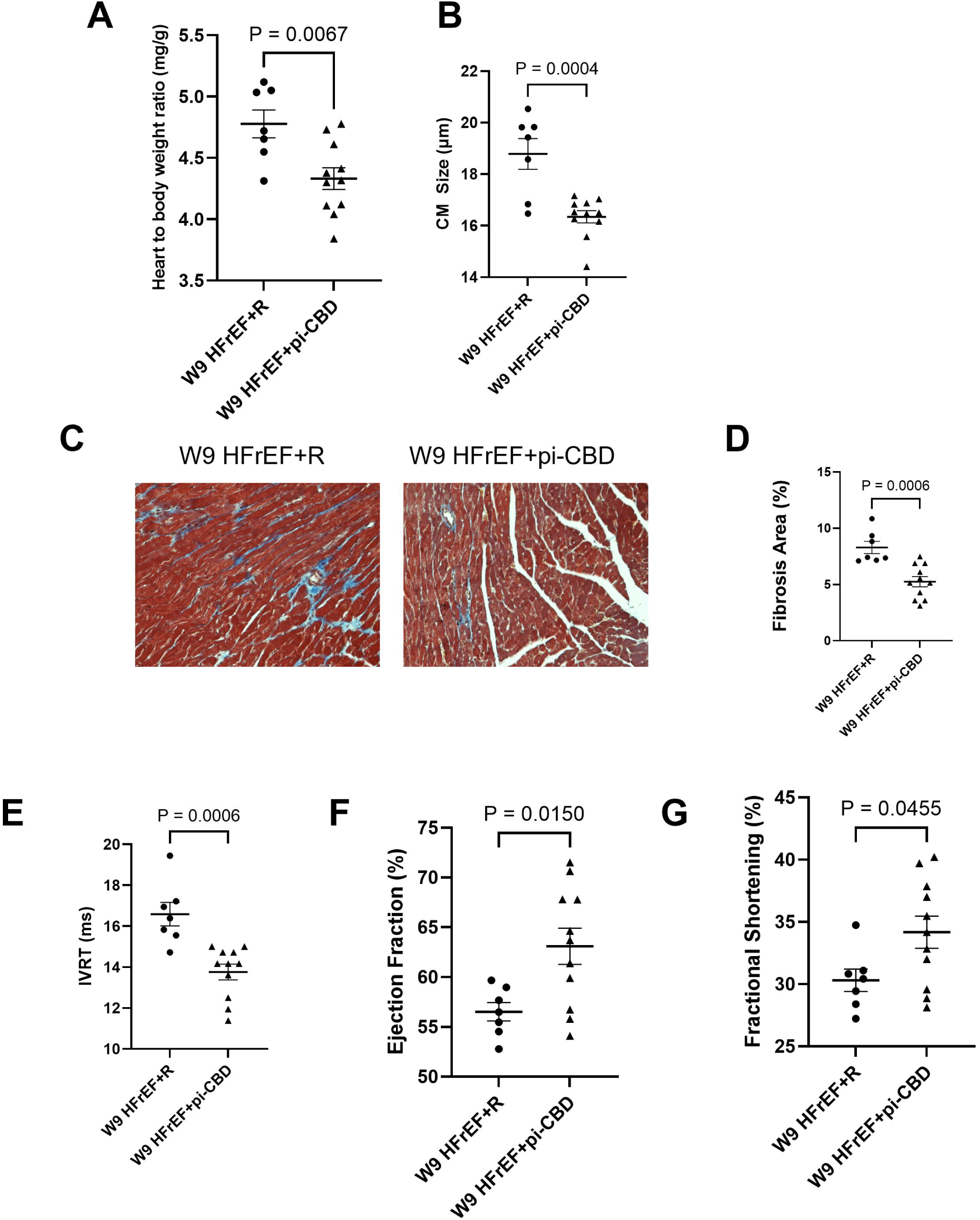
Post-induction synthetic cannabidiol treatment attenuates residual adverse remodeling and improves diastolic function during recovery from experimental HFrEF. **A**, Heart weight-to-body weight ratio (HW/BW), demonstrating persistent cardiac hypertrophy in W9 HFrEF+R mice and its attenuation with post-induction synthetic cannabidiol treatment. **B**, Quantification of cardiomyocyte size, showing reduced cardiomyocyte hypertrophy with post-induction synthetic cannabidiol treatment compared with natural recovery. **C**, Representative Masson trichrome-stained myocardial sections from W9 HFrEF+R and W9 HFrEF+pi-cannabidiol hearts. Collagen is shown in blue and myocardium in red. Scale bar, 50 μm. **D**, Quantification of myocardial fibrotic area, demonstrating reduced residual collagen deposition with post-induction synthetic cannabidiol treatment compared with natural recovery. **E**, Isovolumetric relaxation time (IVRT), demonstrating improved diastolic relaxation with post-induction synthetic cannabidiol treatment. **F**, Left ventricular ejection fraction (EF), showing substantial systolic recovery in both groups at week 9. **G**, Fractional shortening (FS), similarly demonstrating recovery of systolic performance in both groups at week 9. For **A** and **D– G**, groups were compared using an unpaired 2-tailed Student *t* test. For **B**, groups were compared using the Mann–Whitney *U* test. Data are presented as mean±SEM. *P*<0.05 was considered statistically significant. HFrEF indicates heart failure with reduced ejection fraction; HFrEF+R, HFrEF followed by natural recovery; and HFrEF+pi-cannabidiol, HFrEF treated with synthetic cannabidiol after disease induction; FS, fractional shortening and EF, ejection fraction

### Concomitant s-cannabidiol therapy partially re-aligns the HFrEF transcriptome through a multimodal axis centered around immunofibrotic axis

To define the transcriptional changes accompanying the structural and functional effects of s-cannabidiol, we performed bulk RNA-sequencing on left ventricular tissue from W5 Ctrl, W5 HFrEF, and W5 HFrEF+c-s-cannabidiol mice. To assess whether s-cannabidiol altered the transcriptome in non-diseased myocardium W5 Ctrl+s-cannabidiol mice hearts were also profiled. This group closely resembled W5 CTRL (**Supplementary Figure 3**) and was therefore not included in subsequent disease-focused comparisons. Global transcriptomic analysis demonstrated clear separation among control, HFrEF, and s-cannabidiol-treated HFrEF samples on principal component analysis (**Supplementary Figure 4**), indicating that both disease induction and treatment status contributed to the major axes of transcriptional variation. In these analyses, W5 HFrEF+c-s-cannabidiol samples showed partial re-alignment away from the untreated HFrEF transcriptional profile, without complete convergence with the control state. This pattern supports selective modulation of disease-associated gene-expression programs rather than uniform transcriptomic normalization.

GSEA analysis between W5 HFrEF and W5 HFrEF+c-s-cannabidiol myocardium for the GO terms **(Figure 3A)** identified enrichment of terms related to fatty acid β-oxidation and catabolism, mitochondrial respiratory chain assembly, mitochondrial translation, ribosome biogenesis, cytoplasmic translation, oxidoreductase activity, chromatin remodeling, and receptor kinase activity (**Data File A**). MSigDB Hallmark GSEA identified enrichment of pathways related to fatty acid metabolism, bile acid metabolism, adipogenesis, peroxisome biology, WNT/β-catenin signaling, hedgehog signaling, mitotic spindle, DNA repair, p53 signaling, apoptosis, reactive oxygen species response, oxidative phosphorylation, mTORC1 signaling, TNF-α signaling via NF-κB, hypoxia, unfolded protein response, epithelial–mesenchymal transition, E2F targets, MYC targets, heme metabolism, myogenesis, and estrogen response. (**Figure 3B; Data file B**) Together, these ranked-gene analyses indicate that the morphological normalization seen in the concomitant s-cannabidiol treated mice is associated with selective pathway-level remodeling of metabolic, mitochondrial, translational, stress-response, inflammatory, cell-cycle, and fibrotic remodeling-associated transcriptional programs without global normalization of all disease associated pathways.

**Figure 3.**
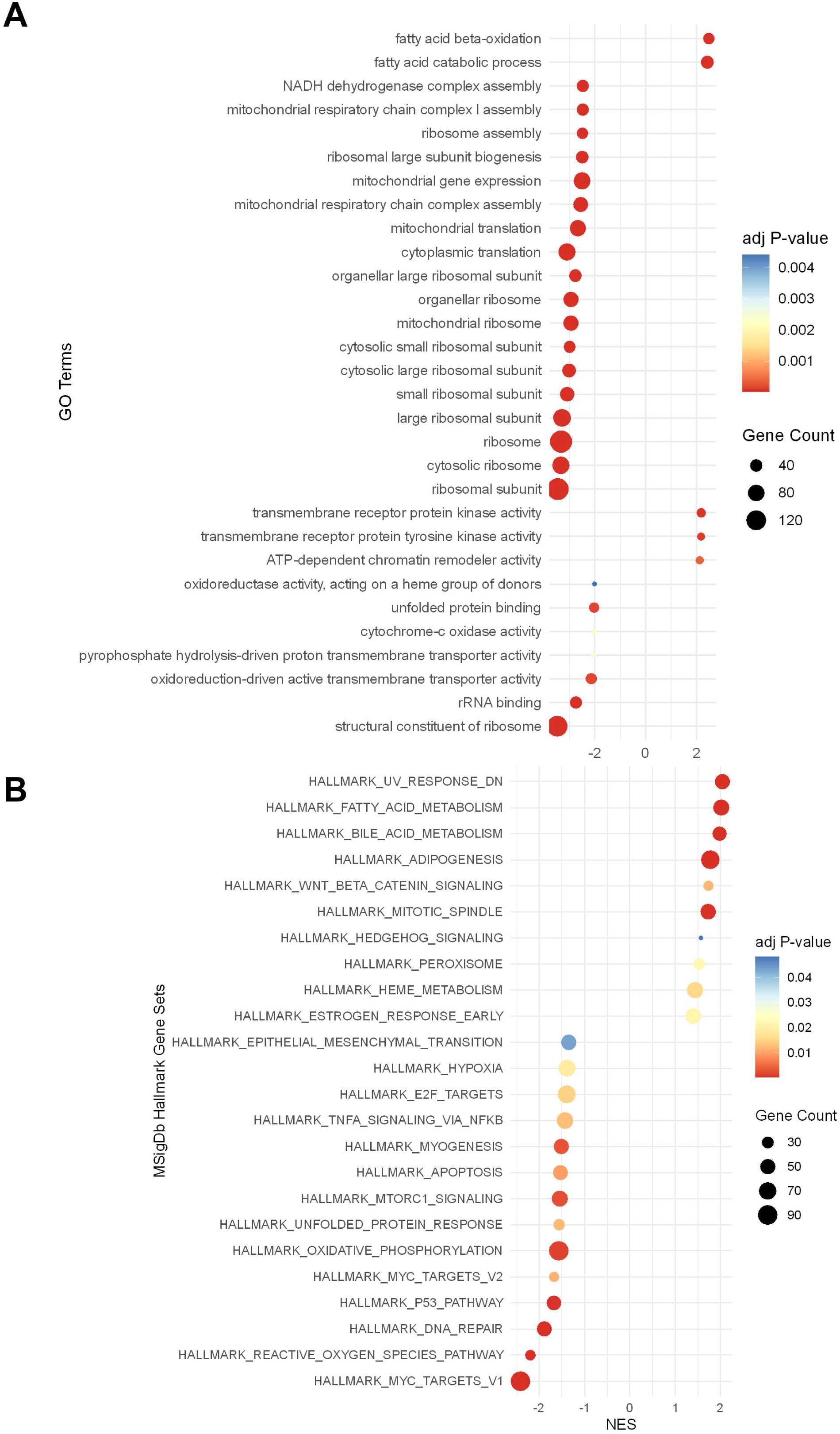
Concomitant synthetic cannabidiol treatment induces selective transcriptional reprogramming of metabolic, mitochondrial, and stress-response pathways in experimental HFrEF. **A**, Gene Ontology (GO) gene set enrichment analysis (GSEA) of ranked genes comparing W5 HFrEF treated concomitantly with synthetic cannabidiol with untreated W5 HFrEF. **B**, Molecular Signatures Database (MSigDB) Hallmark GSEA demonstrating pathway-level changes associated with concomitant synthetic cannabidiol treatment. Positive normalized enrichment scores (NES) indicate pathways enriched with cannabidiol treatment, whereas negative NES values indicate pathways negatively enriched with cannabidiol treatment relative to HFrEF. NES indicates normalized enrichment score. Dot size represents gene count, and color indicates adjusted *P* value. W5 HFrEF, week 5 heart failure with reduced ejection fraction; MSigDb indicates Molecular Signatures Database.

DEG analysis further showed that HFrEF induction was associated with broad bidirectional transcriptional remodeling (**Figure 4 A&B**), whereas the W5 HFrEF versus W5 HFrEF+c-s-cannabidiol comparison identified a smaller subset of cannabidiol-responsive transcripts. In the W5 HFrEF versus W5 Ctrl comparison, 503 genes were upregulated and 473 genes were downregulated, while the W5 HFrEF versus W5 HFrEF+c-s-cannabidiol comparison identified 80 upregulated and 122 downregulated transcripts. Overlap analysis indicated that s-cannabidiol modified a subset of the HFrEF-associated transcriptional program, affecting approximately 12% of genes upregulated in HFrEF relative to control mice and 22% of genes downregulated in HFrEF relative to control (**Figure 4 C &D**). Consistent with this, the scatterplot comparing disease-associated and treatment-associated log2 fold changes showed that most overlapping genes changed in opposite directions between the HFrEF versus CTRL and HFrEF+c-s-cannabidiol versus HFrEF contrasts, supporting broad directional attenuation of the HFrEF transcriptional signature by s-cannabidiol (**Figure 4B**). We next interrogated the biological processes represented within the s-cannabidiol-responsive transcriptome with pathway enrichment performed using only the DEG set identified between W5 HFrEF and W5 HFrEF+c-s-cannabidiol groups. Overrepresentation analysis using the Gene ontology (GO) gene set library of EnrichR showed that the most enriched terms were dominated by plasma membrane, ion/transmembrane transport, regulation of signaling, and transcriptional regulation terms. The nervous system process in the biological process and the neuron projection in the cellular component segment seem to represent genes related to excitable-cell, membrane signaling, ion channel, neuropeptide and guidance/adhesion genes. **(Supplementary Figure 5A, Data File C)** These categories are consistent with published mechanisms of s-cannabidiol cardio protection involving preserved calcium handling, mitochondrial function, redox balance, and anti-inflammatory signaling. Additional enriched categories included cellular response to hormone stimulus, monoatomic cation transmembrane transport, and multiple axon guidance/projection-related processes. Molecular function enrichment further highlighted sodium channel activity, voltage-gated sodium channel activity, cyclic nucleotide binding, and cyclic nucleotide-activated cation channel activity, indicating that the differential transcriptome also included membrane signaling and transport-related programs. These terms appear to reflect s-cannabidiols counter to selected nodes of HF pathology, in contrast to the broad HF signature GO terms seen in **Supplementary Figure 5** (showing an impact on metabolic remodeling, oxidative/redox biology, cell-cycle proliferative remodeling, Wnt/TGF-B like signal, contractile/sarcomere remodeling and ECM/fibrosis). Overrepresentation analysis exploring the Hallmark MSigDb gene set library yielded only one statistically significant term, namely Epithelial Mesenchymal Transition (EMT) **(Supplementary Figure 4B, Data file D)**, suggesting that the most prominent higher-order pathway signal within the s-cannabidiol-responsive transcriptome is linked to fibrotic remodeling mediated through non-cardiomyocyte phenotype transitions. A gene level grouping of all the DEGs into biological modules and comparison to published literature is presented in **Supplementary Table 1** as hypothesis generating for the possible impact of s-cannabidiol in treating heart failure.

**Figure 4.**
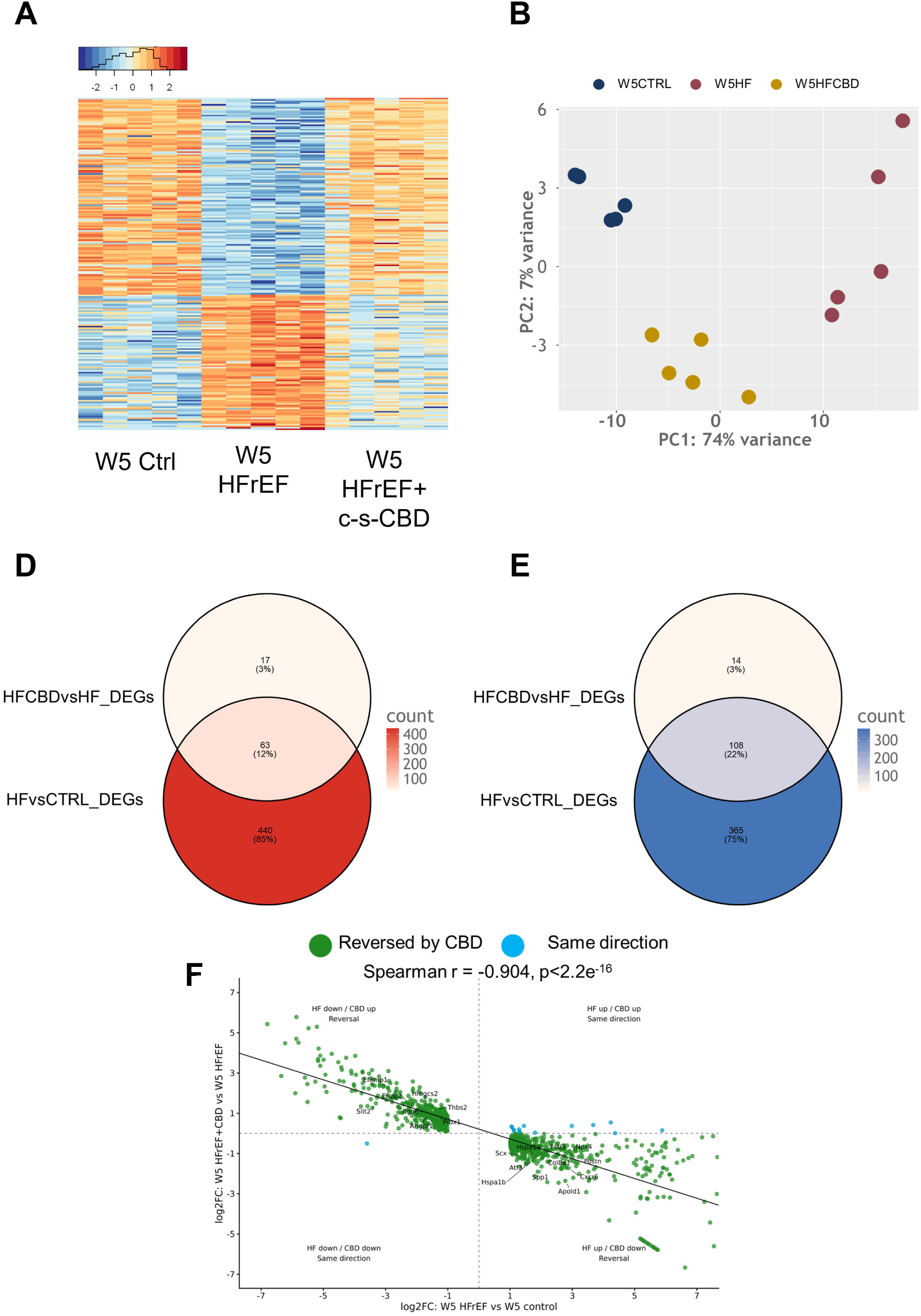
Concomitant synthetic cannabidiol treatment selectively remodels and directionally opposes the HFrEF-associated transcriptional program. **A**, Heatmap of differentially expressed genes (DEGs) across W5 CTRL, W5 HFrEF, and W5 HFrEF treated concomitantly with synthetic cannabidiol. W5 HFrEF samples exhibit a distinct transcriptional profile compared with W5 CTRL, whereas cannabidiol-treated HFrEF samples show a marked shift away from the untreated HFrEF expression pattern. Each column represents an individual animal, and each row represents a gene. Relative gene expression patterns are displayed from lower expression in blue to higher expression in red. **B**, Principal component analysis (PCA) of global gene expression demonstrating separation of W5 HFrEF from W5 CTRL samples. Cannabidiol-treated HFrEF samples form a distinct cluster displaced from the untreated HFrEF group, consistent with treatment-associated transcriptional remodeling. **D–E**, Overlap analyses comparing DEGs identified in the W5 HFrEF versus W5 CTRL contrast with genes differentially expressed in response to concomitant cannabidiol treatment, illustrating the subset of disease-associated transcriptional changes affected by cannabidiol. **F**, Scatterplot comparing the disease-associated log2 fold change (W5 HFrEF versus W5 CTRL; x axis) with the cannabidiol-associated log2 fold change (W5 HFrEF+c-s-cannabidiol versus W5 HFrEF; y axis) for HFrEF-associated DEGs defined by FDR adjusted P<0.05 and |log2 fold change|≥1. Genes with missing log2 fold-change values in the s-cannabidiol contrast were excluded; statistical significance in the s-cannabidiol contrast was not required. Each point represents an individual gene. Genes in the upper-left and lower-right quadrants changed in opposite directions in the disease and cannabidiol-treatment contrasts and were classified as directionally reversed by cannabidiol, whereas genes in the upper-right and lower-left quadrants changed in the same direction. Of 972 HFrEF-associated DEGs, 957 genes (98.5%) changed in the opposite direction following cannabidiol treatment, whereas 15 genes (1.5%) changed in the same direction. Disease-associated and cannabidiol-associated transcriptional changes were strongly inversely correlated (Spearman *r*=−0.904, *P*<2.2×10^−16), demonstrating broad directional opposition of the HFrEF-associated transcriptional program by cannabidiol. CTRL indicates control; DEG, differentially expressed gene; HFrEF, heart failure with reduced ejection fraction; and PCA, principal component analysis.

Because GSEA and DEG highlighted matrix remodeling and epithelial– mesenchymal transition–related programs as one of the relevant mechanisms, we next examined curated gene panels relevant to EndMT, endothelial phenotype, mesenchymal phenotype, fibrosis, inflammation, and inflammasome signaling. These panels included 35 EndMT-associated genes, 14 endothelial phenotype-associated genes, 13 mesenchymal phenotype-associated genes (**Figure 5**), 31 fibrosis-associated genes, 23 inflammation-associated genes, and 15 inflammasome-related genes (**Supplementary Figure 6**). Heat maps were generated from bulk RNA-seq count-derived expression values and displayed as row-wise z scores across W5 CTRL, W5 HFrEF, and W5 HFrEF+c-s-cannabidiol samples. Because these heat maps were generated from curated biologically relevant gene sets rather than unbiased lists, they are presented as descriptive, hypothesis-generating visualizations of pathway-related expression patterns. Genes meeting the predefined criteria of absolute log2 fold change ≥0.5 and adjusted *P*<0.05 are marked with an asterisk. A complete gene-level description with statistics are provided in **Supplementary Table 2** and **Data File E**. Within this framework, the focused heat maps suggest selective modulation of endothelial/mesenchymal plasticity, extracellular matrix remodeling, inflammatory signaling, and inflammasome-associated programs in W5 HFrEF+c-s-cannabidiol myocardium. However, these gene level analyses should be considered exploratory and hypothesis generating.

**Figure 5.**
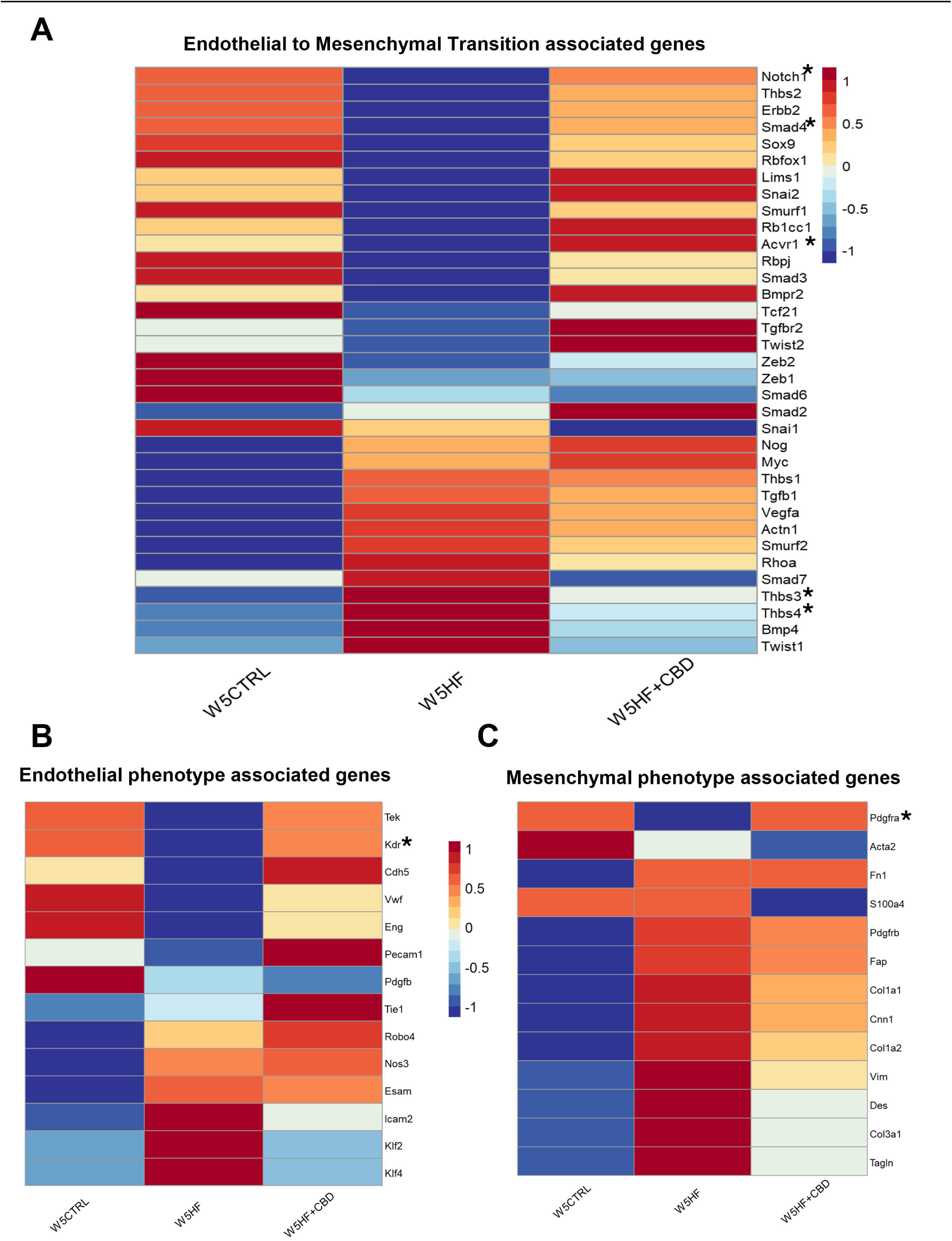
Concomitant synthetic cannabidiol treatment modulates endothelial-to-mesenchymal transition and vascular remodeling–associated transcriptional signatures in experimental HFrEF. **A–C**, Focused analysis of curated gene sets selected on the basis of their established roles in endothelial-to-mesenchymal transition (EndMT), endothelial homeostasis, and mesenchymal activation. Heatmaps compare W5 CTRL, W5 HFrEF, and W5 HFrEF treated concomitantly with synthetic cannabidiol. **A**, EndMT-associated gene signature. HFrEF is associated with coordinated alterations in genes linked to endothelial phenotypic transition, whereas concomitant cannabidiol treatment shifts the expression pattern of this signature relative to untreated HFrEF. **B**, Endothelial function–associated gene signature, comprising genes involved in endothelial identity, vascular homeostasis, nitric oxide signaling, and barrier integrity. Cannabidiol treatment is associated with preservation or restoration of endothelial-associated transcriptional patterns relative to untreated HFrEF. **C**, Mesenchymal and fibroblast activation–associated gene signature, including genes involved in mesenchymal identity, cytoskeletal remodeling, fibroblast activation, and extracellular matrix regulation. HFrEF-associated changes in this gene set are attenuated or directionally remodeled with concomitant cannabidiol treatment. Gene expression values were row-scaled and expressed as *z* scores to visualize relative expression differences among experimental groups. Red indicates relatively higher expression, and blue indicates relatively lower expression within each gene. Columns represent experimental groups, and rows represent individual genes. CTRL indicates control; EndMT, endothelial-to-mesenchymal transition; and HFrEF, heart failure with reduced ejection fraction.

### Post-induction/injury s-cannabidiol therapy maintains therapeutic efficacy and promotes selective transcriptional remodeling beyond natural recovery

Having established that concomitant s-cannabidiol treatment modulated global and pathway-specific transcriptomic programs during HFrEF induction, we next examined whether s-cannabidiol administered after onset of injury was associated with transcriptional changes compared to HFrEF state. As a contextual disease-stage comparison, we first compared transcriptomic profiles from W5 HFrEF and W9 HFrEF+pi-s-cannabidiol mice, recognizing that this comparison reflects the combined influence of time, withdrawal of HFrEF-inducing stimuli, recovery duration, and treatment. This analysis **(Supplementary Figure 8)** showed substantial transcriptional differences between untreated W5 HFrEF and W9 HFrEF+pi-s-cannabidiol myocardium, including distinct heat-map clustering, broad differential gene expression on volcano plot analysis, and enrichment of pathways related to extracellular matrix organization, cell migration, cytoskeletal dynamics, ion channel activity, membrane-associated processes, epithelial–mesenchymal transition, inflammatory signaling, and cell-cycle regulation. Because this comparison is not a clean treatment-only contrast, these findings are best interpreted as evidence that the late-treatment recovery state is transcriptionally distinct from the established W5 HFrEF state rather than as definitive proof of treatment-driven molecular normalization.

Ranked-gene GSEA comparing W9 HFrEF+Recovery and W9 HFrEF+pi-s-cannabidiol identified enrichment of GO terms **(Figure 6A, Data File F)** related to histone methylation, chromatin regulation, mitochondrial and ribosomal pathways, collagen-containing extracellular matrix, complement binding, oxidoreductase activity, and ribosomal structural constituents. MSigDB Hallmark GSEA showed **(Figure 6B, Data File G)** enrichment of pathways including DNA repair, complement, apoptosis, oxidative phosphorylation, xenobiotic metabolism, mTORC1 signaling, E2F targets, coagulation, epithelial–mesenchymal transition, and MYC targets V1. Together, these analyses indicate that post-induction s-cannabidiol treatment is associated with selective transcriptional remodeling during recovery from HFrEF, involving pathways related to epigenetic regulation, metabolism, protein synthesis, extracellular matrix remodeling, immune signaling, cell-cycle regulation, and stress-response programs. Importantly, these data suggest a distinct s-cannabidiol-associated recovery-phase transcriptional state relative to natural recovery as reflected in further improvement of the histological characteristics in the treated mice when compared to natural recovery.

**Figure 6.**
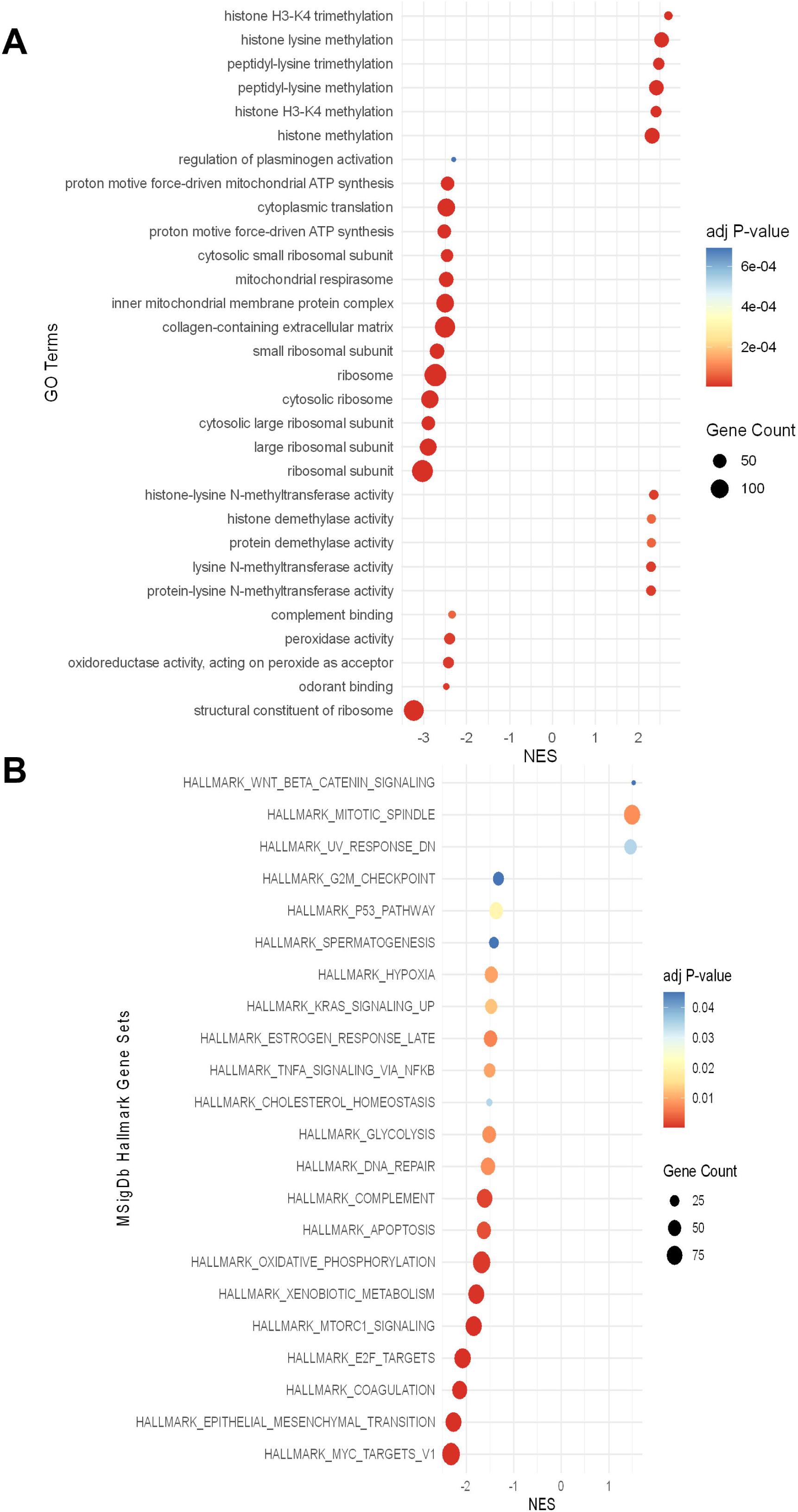
Post-induction synthetic cannabidiol treatment remodels myocardial transcriptional programs during recovery from experimental HFrEF. **A**, Gene set enrichment analysis (GSEA) of Gene Ontology (GO) terms comparing myocardial transcriptomes from W9 HFrEF+pi-cannabidiol and W9 HFrEF+R mice. The dot plot displays significantly enriched GO terms according to normalized enrichment score (NES). Positive NES values indicate enrichment in the post-induction cannabidiol-treated group relative to natural recovery, whereas negative NES values indicate enrichment in the natural recovery group relative to cannabidiol treatment. **B**, GSEA of Molecular Signatures Database (MSigDB) Hallmark gene sets comparing W9 HFrEF+pi-cannabidiol with W9 HFrEF+R myocardium. Positive NES values indicate relative enrichment with post-induction cannabidiol treatment, whereas negative NES values indicate relative enrichment in the natural recovery group. In both (A&B), Dot size represents gene count, and color represents the adjusted *P* value. GO, Gene Ontology; GSEA, gene set enrichment analysis; HFrEF, heart failure with reduced ejection fraction; MSigDB, Molecular Signatures Database; NES, normalized enrichment score; PCA, principal component analysis; HFrEF+R, HFrEF followed by natural recovery; and HFrEF+pi-cannabidiol, HFrEF treated with synthetic cannabidiol after disease establishment.

### Cannabidiol inhibits EndMT in vitro through CB2- and PPARγ-dependent signaling

These in vivo transcriptomic findings prompted us to test directly whether s-cannabidiol modulates endothelial-to-mesenchymal transition in a controlled in vitro model. Using a previously established HUVEC EndMT system, cells were exposed to Ang II/L-NAME, the same stressors used for HFrEF induction in vivo **(Figure 7A)**. Under these conditions, immunofluorescence analysis showed reduced expression of the endothelial marker CD31 together with increased expression of the mesenchymal marker Vimentin, consistent with induction of EndMT **(Figure 7B)**. Concomitant s-cannabidiol treatment suppressed these changes, restoring CD31 and reducing Vimentin expression, thereby preserving endothelial characteristics under pro-EndMT stress. The dose of s-cannabidiol was determined invitro using cell viability **(Supplementary Figure 9)**

**Figure 7.**
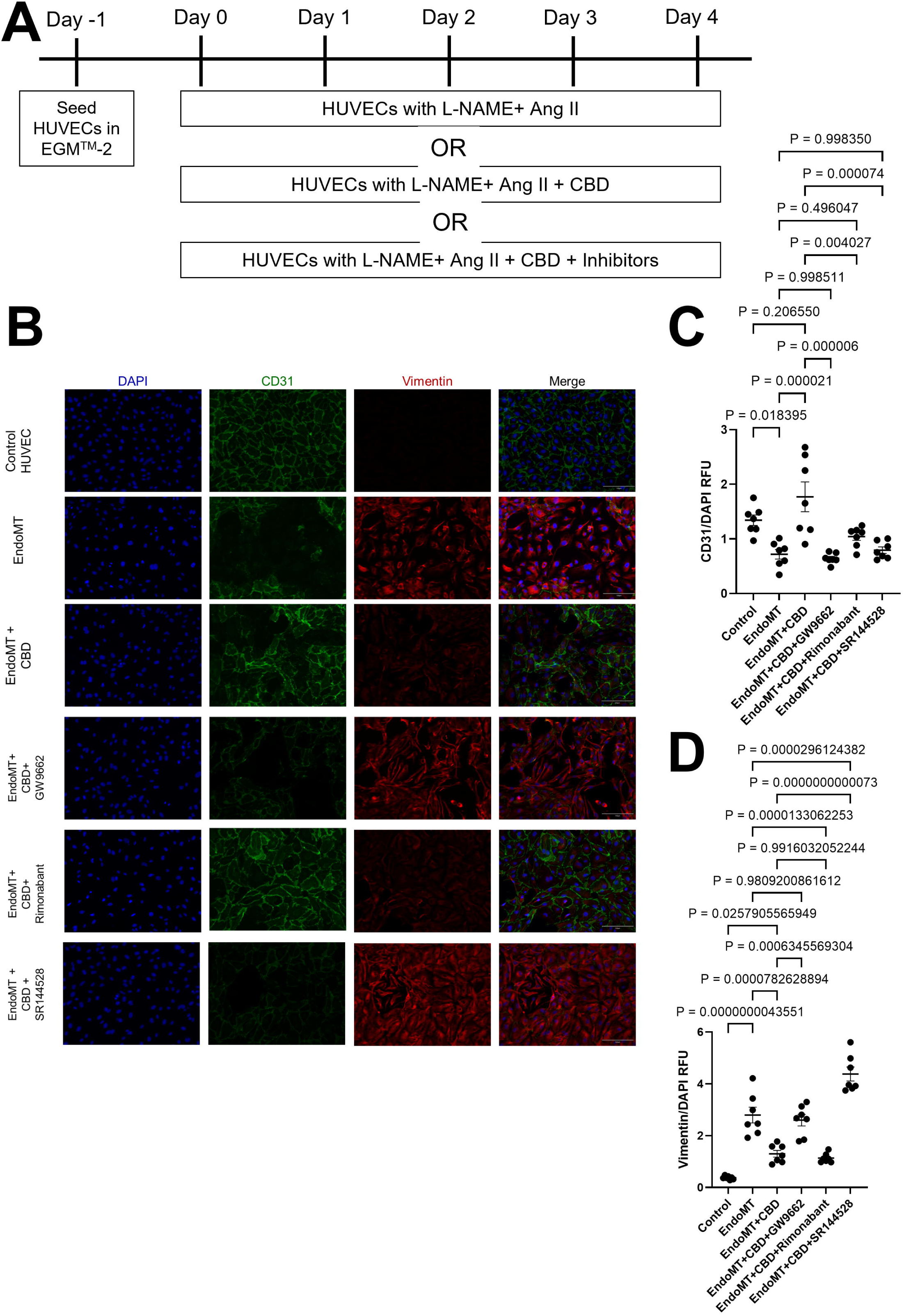
Synthetic cannabidiol suppresses Ang II/L-NAME–induced endothelial-to-mesenchymal transition through CB2- and PPARγ-dependent mechanisms in HUVECs. **A**, Schematic of the in vitro experimental design. Human umbilical vein endothelial cells (HUVECs) were seeded in EGM-2 on day −1 and subsequently exposed for 4 days to Ang II and L-NAME to induce endothelial-to-mesenchymal transition (EndMT). Cells were treated with Ang II+L-NAME alone, Ang II+L-NAME with synthetic cannabidiol, or Ang II+L-NAME with synthetic cannabidiol in the presence of pharmacological inhibitors of PPARγ, CB1, or CB2. **B**, Representative immunofluorescence images of HUVECs stained for the endothelial marker CD31 (green), the mesenchymal marker vimentin (red), and nuclei with DAPI (blue). Experimental groups included untreated control HUVECs, EndMT, EndMT+cannabidiol, EndMT+cannabidiol+GW9662, EndMT+cannabidiol+rimonabant, and EndMT+cannabidiol+SR144528. Scale bar, 150 μm. **C**, Quantification of CD31 fluorescence intensity normalized to DAPI. Cannabidiol increased CD31 signal relative to the EndMT condition. Data were analyzed using 1-way ANOVA followed by Tukey multiple-comparisons test. **D**, Quantification of vimentin fluorescence intensity normalized to DAPI. Data were analyzed using 1-way ANOVA followed by Tukey multiple-comparisons test. Data are presented as mean±SEM. *P*<0.05 was considered statistically significant. Ang II indicates angiotensin II; CB1, cannabinoid receptor type 1; CB2, cannabinoid receptor type 2; DAPI, 4′,6-diamidino-2-phenylindole; EGM-2, Endothelial Cell Growth Medium-2; EndMT, endothelial-to-mesenchymal transition; HUVEC, human umbilical vein endothelial cell; L-NAME, Nω-nitro-L-arginine methyl ester; and PPARγ, peroxisome proliferator-activated receptor γ.

To examine the receptor pathways involved, we used pharmacologic inhibitors targeting PPARγ (GW9662), CB1 (rimonabant), and CB2 (SR144528). Inhibition of PPARγ reversed the protective effect of s-cannabidiol, as shown by reduced CD31 expression and increased Vimentin relative to s-cannabidiol treatment alone **(Figure 7)**. Similarly, CB2 blockade abrogated the anti-EndMT effect of s-cannabidiol, with loss of endothelial marker expression and re-emergence of mesenchymal marker accumulation. In contrast, CB1 inhibition did not materially alter s-cannabidiol mediated restoration of CD31 or suppression of Vimentin. These findings indicate that s-cannabidiol inhibits Ang II/L-NAME–induced EndMT through a signaling axis involving CB2 and PPARγ receptors.

### S-cannabidiol reverses established EndMT-associated phenotypic changes in HUVECs

Having shown that concomitant s-cannabidiol suppresses EndMT and that this effect depends on CB2/PPARγ signaling, we next asked whether s-cannabidiol could also reverse an already established EndMT phenotype akin to our HFrEF+pi-s-cannabidiol in vivo experiment. To address this, HUVECs were first subjected to EndMT-inducing conditions (Day 0-4) and then treated with s-cannabidiol (starting Day 4 – day 8) before immunofluorescence analysis **(Figure 8A)**. **Figure 8 B** shows representative images on Day 4 of EndMT induction associated with reduced CD31 staining, increased Vimentin staining, and a more elongated, spindle-like morphology relative to endothelial control conditions. In contrast, s-cannabidiol treatment after induction and assessment on Day 8 showed a relative increase in CD31 signal, reduced Vimentin staining, and preservation of endothelial morphology. Quantitative analysis supported these observations: CD31 fluorescence intensity was significantly reduced by EndMT induction and significantly increased following s-cannabidiol treatment **(Figure 8C)**, whereas Vimentin fluorescence intensity was elevated under EndMT-inducing conditions and significantly decreased after s-cannabidiol administration **(Figure 8D)**. Taken together, these data indicate that s-cannabidiol not only inhibits EndMT induction but can also reverse established EndMT-associated phenotypic changes in vitro.

**Figure 8.**
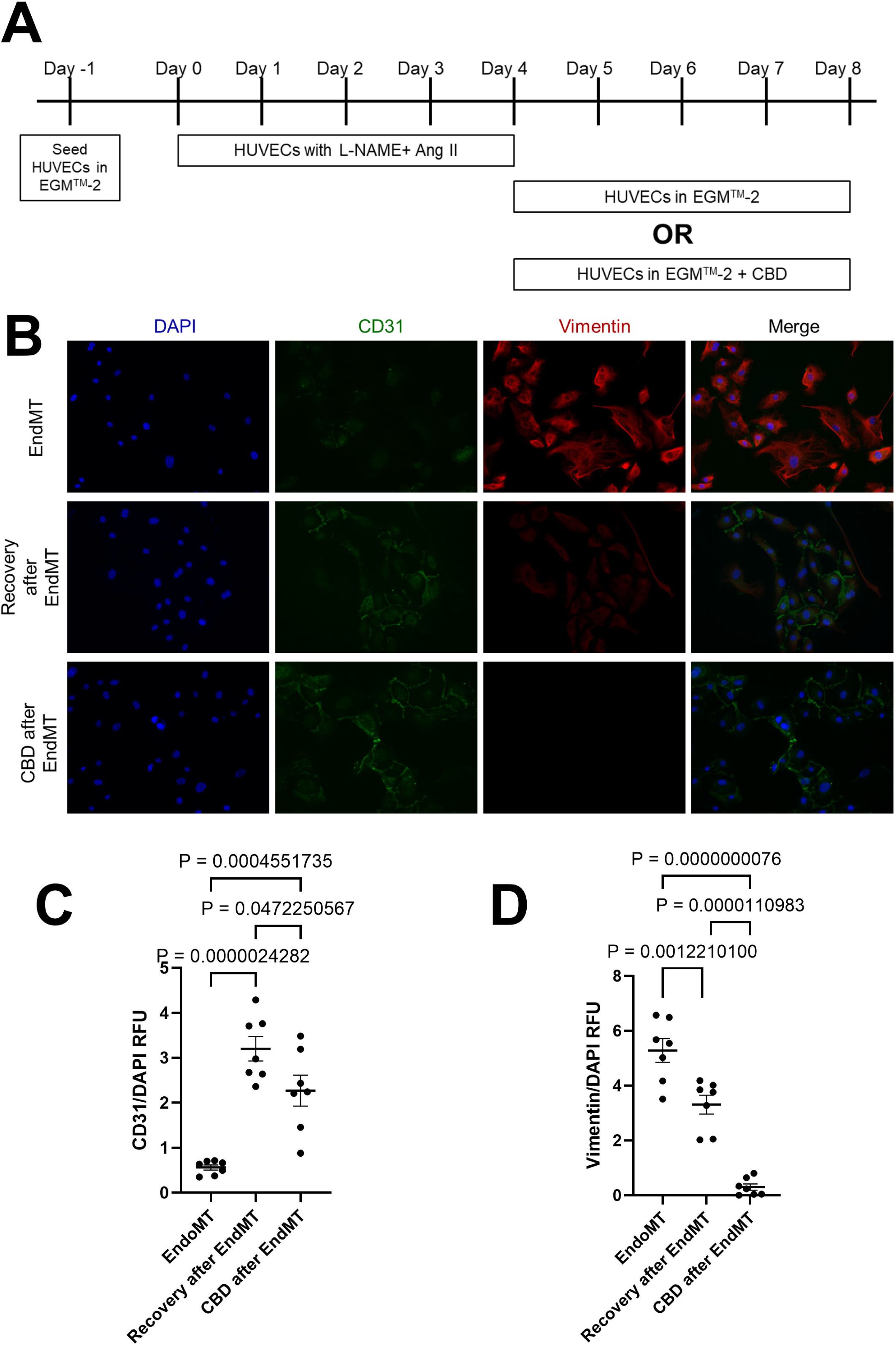
Post-induction synthetic cannabidiol treatment promotes phenotypic recovery from an established endothelial-to-mesenchymal transition–like phenotype in HUVECs. **A**, Schematic of the in vitro recovery experimental design. Human umbilical vein endothelial cells (HUVECs) were seeded in EGM-2 on day −1 and exposed to L-NAME and Ang II from days 0 through 4 to induce an endothelial-to-mesenchymal transition (EndMT)-like phenotype. Following EndMT induction, the inducing stimuli were withdrawn, and cells were cultured from days 4 through 8 in EGM-2 alone to assess natural recovery or in EGM-2 supplemented with synthetic cannabidiol to evaluate treatment-assisted phenotypic recovery. **B**, Representative immunofluorescence images of HUVECs stained for the endothelial marker CD31 (green), the mesenchymal marker vimentin (red), and nuclei with DAPI (blue). Cells exposed to EndMT-inducing conditions exhibited reduced CD31 expression, increased vimentin expression, and an elongated, spindle-like morphology. During recovery after withdrawal of the EndMT-inducing stimuli, synthetic cannabidiol treatment was associated with increased CD31 expression, reduced vimentin expression, and partial restoration of an endothelial-like morphology compared with natural recovery alone. Scale bar, 150 μm. **C**, Quantification of CD31 fluorescence intensity normalized to DAPI. Synthetic cannabidiol treatment during the recovery phase increased CD31 signal compared with the EndMT condition and natural recovery alone. Data were analyzed using 1-way ANOVA followed by Tukey multiple-comparisons test. **D**, Quantification of vimentin fluorescence intensity normalized to DAPI. EndMT induction increased vimentin expression, whereas synthetic cannabidiol treatment during the recovery phase reduced vimentin signal relative to the EndMT condition and natural recovery alone. Data were analyzed using 1-way ANOVA followed by Tukey multiple-comparisons test. Data are presented as mean±SEM. *P*<0.05 was considered statistically significant. Ang II indicates angiotensin II; DAPI, 4′,6-diamidino-2-phenylindole; EGM-2, Endothelial Cell Growth Medium-2; EndMT, endothelial-to-mesenchymal transition; HUVEC, human umbilical vein endothelial cell; and L-NAME, Nω-nitro-L-arginine methyl ester.

## Discussion

HFrEF is characterized not only by impaired contractile performance but also by persistent fibrotic remodeling, altered interstitial architecture, endothelial dysfunction, and inflammatory activation. ^11, 12^ The contribution of the microvascular endothelium, interstitial milieu, and local tissue inflammation to biological remodeling in heart failure continues to evolve. ^2^ Cannabidiol has emerged as a candidate cardioprotective compound because of its reported anti-inflammatory and anti-fibrotic properties, yet the mechanisms by which cannabidiol may modulate cardiac remodeling in heart failure remain incompletely defined. ^5, 6^ Because naturally extracted cannabidiol preparations may vary in composition, we evaluated a synthetic, pharmaceutically manufactured cannabidiol formulation. In this study, s-cannabidiol attenuated the development of structural and functional remodeling when administered concomitantly with HFrEF induction and also showed therapeutic benefit when administered after injury onset. Importantly, in the post induction/injury recovery model, s-cannabidiol further improved residual structural and transcriptional remodeling despite substantial spontaneous recovery of systolic indices, suggesting that improvement in ejection fraction alone may not fully capture ongoing tissue-level pathology.

Across the in vivo models, our data supports a framework in which s-cannabidiol favorably modulates cardiac remodeling through coordinated effects on extracellular matrix, inflammatory signaling, endothelial/mesenchymal plasticity, and stress-adaptive transcriptional programs. The translational relevance of this product is supported by a recent international clinical trial, suggesting (safety and) potential benefit of a pharmaceutically produced s-cannabidiol in acute myocarditis where cardiac magnetic resonance imaging changes showed a significant reduction in left ventricular mass in treated patients compared to placebo. ^13^ While acute myocarditis is a different entity than other HFrEF etiologies, there remains a commonality of immune activation and response, albeit to different degrees in these two clinical entities.

In clinical practice, therapeutic interventions are rarely initiated at the earliest onset of molecular injury, and many patients are treated after substantial remodeling has already occurred. ^14^ By using a experimental design in which treatment started after HFrEF-inducing stimuli were established, we were able to examine whether s-cannabidiol provides benefit despite onset of substantial remodeling. Another key implication of our recovery after HFrEF induction model is that organ level functional recovery, structural recovery, and molecular recovery are not synonymous. ^15, 16^ Although systolic indices improved substantially in the natural recovery group, residual hypertrophy and fibrosis persisted, and s-cannabidiol further improved these structural endpoints. These findings emphasize that normalization or improvement of echocardiographic indices does not necessarily imply complete restoration of the underlying biological state. ^15, 16^ This concept is also supported by the transcriptomic analyses, which showed that s-cannabidiol modified disease-associated transcriptional programs further, when compared to natural/passive recovery.

The concomitant-treatment transcriptomic analyses indicate that s-cannabidiol partially re-aligns the HFrEF transcriptome rather than normalization to a control state. Heat-map clustering, PCA, DEG analysis, fold-change scatterplots, and overlap analyses showed that HFrEF induction produced broad bidirectional transcriptional remodeling, while s-cannabidiol modified a smaller subset of the HFrEF-associated transcriptional program that was sufficient to create a morphological preservation. The observation that many overlapping genes changed in opposite directions between the disease and treatment state supports directional attenuation of the HFrEF transcriptional signature. Hence, this should be interpreted as partial and selective transcriptomic re-alignment rather than complete molecular normalization. This distinction is important because substantial structural and functional benefit may occur even when the myocardium retains a residual molecular imprint of prior injury ^15, 16^. Whether higher doses, longer treatment duration, or alternative therapeutic windows would produce more complete transcriptomic recovery remains to be determined.

The GSEA analyses provides a broader view of the biological domains associated with s-cannabidiol treatment. In the W5 HFrEF versus W5 HFrEF+c-s-cannabidiol comparison, GO-based GSEA identified enrichment of pathways related to fatty acid β-oxidation and catabolism, mitochondrial respiratory chain assembly, mitochondrial translation, ribosome biogenesis, cytoplasmic translation, oxidoreductase activity, chromatin remodeling, and receptor kinase activity. MSigDB Hallmark GSEA further identified enrichment of metabolic, mitochondrial, stress-response, inflammatory, cell-cycle, DNA repair, MYC/E2F, mTORC1, epithelial–mesenchymal transition, and remodeling-associated pathways. These findings suggest that s-cannabidiol treatment is associated with broad pathway-level remodeling involving cardiac metabolism, mitochondrial function, protein synthesis, stress-response signaling, inflammatory activation, and extracellular matrix remodeling. Importantly, these pathway-level results should not be interpreted as evidence that every enriched program is uniformly normalized, but rather as evidence that s-cannabidiol reshapes multiple biological networks activated or suppressed during the injury sustained through HFrEF induction.

The focused curated gene assessment provides a hypothesis generation. Within that framework, HFrEF was associated with altered expression patterns across curated EndMT-associated, endothelial phenotype-associated, mesenchymal phenotype-associated, fibrosis-associated, inflammation-associated, and inflammasome-related genes. S-cannabidiol treatment shifted several of these curated signatures in a direction consistent with reduced EndMT, pro endothelial and reduction of mesenchymal state in a reduced inflammatory milieu. These data cannot establish cell-type-specific transcriptional dynamics or prove endothelial-to-mesenchymal transition in vivo as it reflects bulk RNA-seq alone.^17^

The post-induction recovery transcriptomic analyses extend these findings. The comparison between W5 HFrEF and W9 HFrEF+pi-s-cannabidiol demonstrated marked transcriptional differences; however, this comparison reflects the combined effects of time, withdrawal of HFrEF-inducing stimuli, recovery duration, and treatment. The more direct comparison between W9 HFrEF+Recovery and W9 HFrEF+pi-s-cannabidiol showed that s-cannabidiol-treated hearts occupied a distinct recovery-phase transcriptional state. Ranked-gene GSEA in this comparison identified GO terms related to histone methylation, chromatin regulation, mitochondrial and ribosomal pathways, collagen-containing extracellular matrix, complement binding, oxidoreductase activity, and ribosomal structural constituents. Hallmark GSEA identified pathways including DNA repair, complement, apoptosis, oxidative phosphorylation, xenobiotic metabolism, mTORC1 signaling, E2F targets, coagulation, epithelial–mesenchymal transition, and MYC targets V1. Together, these analyses suggest that post-induction s-cannabidiol treatment is associated with selective transcriptional remodeling during recovery, involving epigenetic regulation, metabolism, protein synthesis, extracellular matrix remodeling, immune signaling, cell-cycle regulation, and stress-response programs. These findings should be interpreted as evidence of a distinct s-cannabidiol-associated recovery-phase molecular state relative to natural recovery or control myocardium in spite of similar (normal range) echocardiographic characteristics in all the three groups.

The in vitro studies provide a mechanistic cellular correlate to the in vivo transcriptomic findings. In HUVECs exposed to Ang II/L-NAME, s-cannabidiol inhibited the acquisition of an EndMT-like phenotype, preserving endothelial marker expression and reducing mesenchymal marker induction. Pharmacologic receptor-blocking studies showed that the anti-EndMT effects of s-cannabidiol were mediated through CB2 and PPARγ receptors. A further translationally relevant observation was that s-cannabidiol not only limited EndMT induction but also partially reversed an already established EndMT-like phenotype after endothelial cells had begun to acquire mesenchymal characteristics. This distinction between prevention and reversal parallels the two therapeutic settings modeled in vivo: treatment during active injury and treatment after injury onset. However, because the in vitro model uses HUVECs and the in vivo transcriptomics were generated from bulk cardiac tissue, these findings should be interpreted as supporting endothelial plasticity as one component of the s-cannabidiol response rather than proving that EndMT is the sole causal mechanism in vivo ^17^.

The present findings also complement prior work suggesting that cannabidiol can modulate cardiomyocyte biology, including pathways related to mitochondrial function and calcium handling. ^6^ Viewed together, the data suggests that s-cannabidiol may exert cardioprotective effects at multiple levels of cardiac organization, influencing cardiomyocyte stress responses, endothelial phenotype preservation, interstitial remodeling, extracellular matrix regulation, and inflammatory signaling. This broader multicellular perspective is particularly relevant in heart failure, where non-myocyte populations and their cross-talk with cardiomyocytes contribute substantially to disease progression and recovery ^2^.

Several limitations should be acknowledged. First, this study examined one s-cannabidiol dose, and dose-response relationships remain to be defined. Second, the work was performed in preclinical models, and the extent to which these findings translate to human HFrEF requires further study. Third, the in vivo transcriptomic analyses were performed in bulk cardiac tissue; therefore, the data implicating cell specific changes are hypothesis generating only and cannot establish cell-type-specific transcriptional dynamics. Fourth, the CB2/PPARγ framework was supported by pharmacologic inhibition in vitro, but genetic perturbation studies would strengthen causal inference. Finally, with s-cannabidiols impact reflected to act through multiple complementary mechanisms, EndMT should be viewed as a contributor to the observed biological benefit rather than the sole mechanism.

In summary, this study identifies pharmaceutically manufactured s-cannabidiol as a modulator of adverse remodeling in experimental HFrEF, with efficacy when administered during disease induction and when administered after injury onset. The in vivo and in vitro data support a model in which s-cannabidiol attenuates pathological hypertrophy and fibrosis, partially re-aligns the failing-heart transcriptome, and modulates endothelial/mesenchymal plasticity and remodeling-associated pathways through mechanisms linked, at least in part, to CB2/PPARγ signaling. These findings support further evaluation of s-cannabidiol as a candidate therapy for heart failure to target residual pathobiological mechanisms beyond the neurohormonal mileu. Our work also highlights that endothelial plasticity, extracellular matrix remodeling, inflammatory signaling, and metabolic/stress-response pathways are potentially targetable components of maladaptive cardiac remodeling. Importantly, these findings are specific to the pharmaceutical formulation studied here and should not be extrapolated to plant-derived cannabidiol preparations, phytocannabinoid mixtures, or other cannabis products.

## Supporting information

Supplementary Figures, Tables and Data fiels

## Acknowledgement

We are grateful to Cardiol Therapeutics for the institutional grant and for providing the pharmaceutically manufactured synthetic cannabidiol in kind.

## Disclosure

This work is partly supported by an institutional grant from Cardiol Therapeutics and the pharmaceutically manufactured synthetic cannabidiol was provided in kind by the same company. GTA serves as the chairman of the board of Cardiol Therapeutics and AB served as a consultant on the steering committee of the ARCHER trial (NCT05180240).

## Figure legends

**Supplementary Figure 1. Experimental design of concomitant and post-induction synthetic cannabidiol treatment paradigms in experimental HFrEF.** Schematic overview of 2 complementary experimental paradigms used to evaluate the effects of synthetic cannabidiol in experimental HFrEF. The upper portion of the schematic depicts the **concomitant treatment paradigm**, comprising W5 CTRL, W5 CTRL+c-cannabidiol, W5 HFrEF, and W5 HFrEF+c-cannabidiol groups. In this paradigm, synthetic cannabidiol was administered at 1 mg/kg by twice-weekly subcutaneous injection during HFrEF induction, enabling assessment of its effects on disease development and adverse cardiac remodeling. The lower portion of the schematic depicts the **post-induction treatment paradigm**, comprising W9 CTRL, W9 HFrEF+R, and W9 HFrEF+pi-cannabidiol groups. In this paradigm, HFrEF was first established during the initial induction phase. Following cessation of the HFrEF-inducing stimuli, mice underwent either natural recovery without cannabidiol treatment or post-induction treatment with synthetic cannabidiol at 1 mg/kg by twice-weekly subcutaneous injection during the recovery phase. Together, these complementary experimental paradigms enabled evaluation of synthetic cannabidiol across distinct therapeutic windows: concomitant treatment during HFrEF development and post-induction treatment during recovery after established cardiac dysfunction. CTRL indicates control; HFrEF, heart failure with reduced ejection fraction; R, natural recovery; c-cannabidiol, concomitant cannabidiol treatment; and pi-cannabidiol, post-induction cannabidiol treatment.

**Supplementary Figure 2. Comparison of cardiac remodeling and ventricular function between established HFrEF and post-induction cannabidiol-treated mice. A**, Heart weight-to-body weight ratio (HW/BW), an index of cardiac hypertrophy. HW/BW was lower in W9 HFrEF+pi-cannabidiol mice than in W5 HFrEF mice. Data were analyzed using an unpaired 2-tailed Student *t* test. **B**, Quantification of cardiomyocyte size from histologic sections. Cardiomyocyte size was reduced in W9 HFrEF+pi-cannabidiol mice compared with W5 HFrEF mice. Data were analyzed using the Mann–Whitney *U* test. **C**, Quantification of myocardial fibrotic area from Masson trichrome-stained sections. Fibrotic burden was lower in W9 HFrEF+pi-cannabidiol hearts than in W5 HFrEF hearts, consistent with reduced residual myocardial fibrosis at the post-induction treatment time point. Data were analyzed using an unpaired 2-tailed Student *t* test. **D**, Isovolumetric relaxation time (IVRT), an index of diastolic function. W9 HFrEF+pi-cannabidiol mice exhibited shorter IVRT compared with W5 HFrEF mice, consistent with improved diastolic relaxation. Data were analyzed using an unpaired 2-tailed Student *t* test. **E**, Left ventricular ejection fraction (EF), an index of systolic function. EF was higher in W9 HFrEF+pi-cannabidiol mice than in W5 HFrEF mice, consistent with improved systolic performance at the later treatment time point. Data were analyzed using an unpaired 2-tailed Student *t* test. **F**, Fractional shortening (FS), an additional measure of systolic function. FS was higher in W9 HFrEF+pi-cannabidiol mice than in W5 HFrEF mice, consistent with improved contractile performance. Data were analyzed using an unpaired 2-tailed Student *t* test. Normality was assessed using the Shapiro–Wilk test. Data are presented as mean±SEM. *P*<0.05 was considered statistically significant. EF indicates ejection fraction; FS, fractional shortening; HFrEF, heart failure with reduced ejection fraction; HW/BW, heart weight-to-body weight ratio; IVRT, isovolumetric relaxation time; and pi-cannabidiol, post-induction synthetic cannabidiol treatment.

**Supplementary Figure 3. Synthetic cannabidiol produces limited gene-level transcriptional changes in non-diseased myocardium. A**, Gene set enrichment analysis (GSEA) of Gene Ontology (GO) terms comparing myocardial transcriptomes from W5 CTRL mice and W5 CTRL mice treated concomitantly with synthetic cannabidiol. Differentially enriched GO terms included pathways related to mitochondrial respiration and ATP synthesis, ribosomal organization and translation, nucleotide biosynthesis, protein transport, oxidoreductase activity, and nuclear receptor–associated processes. Dot size represents gene count, and color represents the adjusted *P* value. **B**, GSEA of Molecular Signatures Database (MSigDB) Hallmark gene sets comparing W5 CTRL and W5 CTRL+c-cannabidiol myocardium. Differentially enriched gene sets included protein secretion, mitotic spindle regulation, UV response, G2M checkpoint, heme metabolism, MYC targets, reactive oxygen species signaling, DNA repair, and oxidative phosphorylation. Dot size represents gene count, and color represents the adjusted *P* value. **C**, Heatmap of differentially expressed genes (DEGs) between W5 CTRL and W5 CTRL+c-cannabidiol myocardium. Each column represents an individual mouse, and each row represents a gene. Gene expression values were row-scaled to visualize relative expression differences between groups. Red indicates relatively higher expression and blue indicates relatively lower expression within each gene. **D**, GO enrichment analysis of DEGs identified between W5 CTRL and W5 CTRL+c-cannabidiol myocardium. Several biological-process terms were represented, including pathways related to immune regulation, apoptotic signaling, RNA metabolic regulation, circadian rhythm, and ATP binding; however, none remained statistically significant after multiple-testing correction. Dot size represents gene count, and color represents statistical significance. Collectively, these analyses indicate that synthetic cannabidiol treatment in non-diseased myocardium is associated with limited gene-level differential expression despite selective pathway-level enrichment. CTRL indicates control; DEG, differentially expressed gene; GO, Gene Ontology; GSEA, gene set enrichment analysis; and MSigDB, Molecular Signatures Database.

**Supplementary Figure 4. Global principal component analysis reveals disease stage– and treatment-associated transcriptomic remodeling.** Principal component analysis (PCA) of global myocardial gene expression across all experimental groups. Each point represents an individual biological sample, and shaded ellipses represent 75% confidence regions. PC1 and PC2 account for 35% and 22% of total transcriptomic variance, respectively. W5 HFrEF samples separate clearly from W5 CTRL samples, consistent with a disease-associated transcriptional shift. W5 HFrEF+c-cannabidiol samples are displaced from the untreated HFrEF cluster, whereas W5 CTRL+cannabidiol samples remain proximal to W5 CTRL samples. At week 9, both W9 HFrEF+R and W9 HFrEF+pi-cannabidiol samples are displaced from the W5 HFrEF transcriptional state and occupy transcriptional spaces partially overlapping with control samples, consistent with spontaneous and treatment-associated transcriptomic remodeling during recovery. CTRL indicates control; HFrEF, heart failure with reduced ejection fraction; PCA, principal component analysis; HFrEF+R, HFrEF followed by natural recovery; c-cannabidiol, concomitant synthetic cannabidiol treatment; and pi-cannabidiol, post-induction synthetic cannabidiol treatment.

**Supplementary Figure 5. Functional enrichment analysis of genes differentially expressed in response to concomitant synthetic cannabidiol treatment in experimental HFrEF. A**, Gene Ontology (GO) enrichment analysis of the complete set of differentially expressed genes (DEGs) identified between W5 HFrEF and W5 HFrEF treated concomitantly with synthetic cannabidiol. Enriched terms encompassed biological processes and molecular functions related to immune regulation, extracellular matrix organization, apoptotic signaling, metabolic regulation, ion channel activity, and other cellular regulatory processes. Dot size represents gene count, and color represents statistical significance. **B**, Molecular Signatures Database (MSigDB) Hallmark pathway enrichment analysis performed using the same DEG set. Epithelial-to-mesenchymal transition was the only Hallmark pathway that remained statistically significant after multiple-testing correction. Other represented pathways did not meet the adjusted significance threshold. Because only a single Hallmark pathway met the predefined significance criterion, a chord plot was not generated for this analysis. DEG indicates differentially expressed gene; GO, Gene Ontology; HFrEF, heart failure with reduced ejection fraction; and MSigDB, Molecular Signatures Database.

**Supplementary Figure 6. Disease-associated transcriptional remodeling and functional enrichment in W5 HFrEF myocardium. A**, Gene set enrichment analysis (GSEA) of Gene Ontology (GO) terms comparing W5 HFrEF and W5 CTRL myocardial transcriptomes. Differentially enriched GO terms encompassed processes related to muscle cell migration, histone acetylation, retinoid homeostasis, lipid and fatty acid metabolism, mitochondrial and ribosomal organization, protein homeostasis, extracellular matrix structure, and oxidoreductase activity. Dot size represents gene count, and color represents the adjusted *P* value. **B**, GSEA of Molecular Signatures Database (MSigDB) Hallmark gene sets comparing W5 HFrEF and W5 CTRL myocardium. Differentially enriched pathways included epithelial–mesenchymal transition, angiogenesis, hypoxia, apoptosis, myogenesis, reactive oxygen species signaling, MYC target programs, mTORC1 signaling, and fatty acid and bile acid metabolism. Dot size represents gene count, and color represents the adjusted *P* value. Positive NES values indicate enrichment in W5 HFrEF relative to W5 CTRL, whereas negative NES values indicate enrichment in W5 CTRL relative to W5 HFrEF. **C**, Heatmap of differentially expressed genes (DEGs) between W5 CTRL and W5 HFrEF myocardium. Each column represents an individual mouse, and each row represents a gene. Gene expression values were row-scaled to visualize relative expression differences among samples. Red indicates relatively higher expression and blue indicates relatively lower expression within each gene. W5 HFrEF samples display a distinct disease-associated transcriptional profile compared with W5 CTRL samples. **D**, GO enrichment analysis of DEGs identified between W5 CTRL and W5 HFrEF myocardium. Represented terms included extracellular matrix organization and structure, angiogenesis, cell migration, cAMP-associated processes, collagen-containing extracellular matrix, receptor–ligand activity, ion channel regulation, and cell–cell adhesion–related functions. Dot size represents gene count, horizontal position represents gene ratio, and color represents the adjusted *P* value. **E**, MSigDB Hallmark pathway enrichment analysis of W5 HFrEF-associated DEGs. Represented pathways included epithelial–mesenchymal transition, G2M checkpoint, E2F targets, TNFα signaling via NF-κB, hypoxia, coagulation, angiogenesis, mitotic spindle, DNA repair, and estrogen response programs. Dot size represents gene count, and color represents statistical significance. Collectively, these analyses demonstrate extensive disease-associated transcriptional remodeling in W5 HFrEF myocardium, involving extracellular matrix remodeling, cellular stress responses, metabolic dysregulation, and proliferative and inflammatory signaling programs. CTRL indicates control; DEG, differentially expressed gene; GO, Gene Ontology; GSEA, gene set enrichment analysis; HFrEF, heart failure with reduced ejection fraction; MSigDB, Molecular Signatures Database; and NES, normalized enrichment score.

**Supplementary Figure 7. Concomitant synthetic cannabidiol treatment remodels fibrosis-, inflammation-, and inflammasome-associated transcriptional signatures in experimental HFrEF.** Focused transcriptomic analysis of curated gene sets related to fibrosis, inflammation, and inflammasome signaling across W5 CTRL, W5 HFrEF, and W5 HFrEF treated concomitantly with synthetic cannabidiol. Heatmaps display relative expression patterns for genes selected on the basis of their established association with the indicated biological processes. Gene expression values were row-scaled and expressed as *z* scores to visualize relative differences among experimental groups. Columns represent experimental groups, and rows represent individual genes. Red indicates relatively higher expression and blue indicates relatively lower expression within each gene. **A**, Fibrosis-associated gene signature comprising genes involved in extracellular matrix organization, collagen synthesis, fibroblast activation, and tissue remodeling. W5 HFrEF myocardium demonstrates coordinated alterations in fibrosis-associated gene expression, whereas concomitant synthetic cannabidiol treatment is associated with attenuation or remodeling of this transcriptional pattern. **B**, Inflammation-associated gene signature comprising cytokines, chemokines, immune signaling mediators, and regulators of inflammatory responses. W5 HFrEF myocardium exhibits an altered inflammatory transcriptional profile, whereas concomitant synthetic cannabidiol treatment is associated with partial attenuation and remodeling of this signature relative to untreated HFrEF. **C**, Inflammasome-associated gene signature comprising genes involved in innate immune activation, inflammasome regulation, and pyroptosis-associated signaling. HFrEF-associated alterations in this gene set are differentially remodeled with concomitant synthetic cannabidiol treatment, consistent with modulation of inflammasome-associated transcriptional programs. CTRL indicates control; HFrEF, heart failure with reduced ejection fraction; and c-cannabidiol, concomitant synthetic cannabidiol treatment.

**Supplementary Figure 8. Comparative transcriptomic profiling of established HFrEF and post-induction cannabidiol-treated myocardium during recovery. A**, Gene set enrichment analysis (GSEA) of Gene Ontology (GO) terms comparing W5 HFrEF myocardium with W9 HFrEF myocardium following post-induction synthetic cannabidiol treatment. Differentially enriched GO terms encompassed fatty acid catabolism and β-oxidation, nucleotide metabolism, tricarboxylic acid cycle–associated processes, cytoskeletal organization, extracellular matrix structure, chemokine binding, and oxidoreductase activity. Dot size represents gene count, and color represents the adjusted *P* value. **B**, GSEA of Molecular Signatures Database (MSigDB) Hallmark gene sets comparing W5 HFrEF with W9 HFrEF+pi-cannabidiol myocardium. Differentially enriched pathways included oxidative phosphorylation, adipogenesis, bile acid and fatty acid metabolism, interferon-α response, TNFα signaling via NF-κB, TGF-β signaling, apoptosis, angiogenesis, and epithelial–mesenchymal transition. Dot size represents gene count, and color represents the adjusted *P* value. Positive NES values indicate enrichment in W9 HFrEF+pi-cannabidiol relative to W5 HFrEF, whereas negative NES values indicate enrichment in W5 HFrEF relative to W9 HFrEF+pi-cannabidiol. **C**, Heatmap of differentially expressed genes (DEGs) between W5 HFrEF and W9 HFrEF+pi-cannabidiol myocardium. Each column represents an individual mouse, and each row represents a gene. Gene expression values were row-scaled to visualize relative expression differences among samples. Red indicates relatively higher expression and blue indicates relatively lower expression within each gene. The 2 groups demonstrate distinct transcriptional profiles, consistent with extensive myocardial transcriptomic remodeling between established HFrEF and the later post-induction cannabidiol-treated recovery state. **D**, GO enrichment analysis of DEGs identified between W5 HFrEF and W9 HFrEF+pi-cannabidiol myocardium. Enriched terms included extracellular matrix organization and structure, regulation of angiogenesis, cell migration and motility, cAMP-associated processes, collagen-containing extracellular matrix, cytoskeletal organization, receptor–ligand activity, ion channel regulation, and cell adhesion–associated functions. Dot size represents gene count, horizontal position represents gene ratio, and color represents the adjusted *P* value. **E**, MSigDB Hallmark pathway enrichment analysis of DEGs identified between W5 HFrEF and W9 HFrEF+pi-cannabidiol myocardium. Represented pathways included epithelial–mesenchymal transition, G2M checkpoint, TNFα signaling via NF-κB, myogenesis, estrogen response, E2F targets, apoptosis, coagulation, KRAS signaling, and angiogenesis. Dot size represents gene count, horizontal position represents gene ratio, and color represents the adjusted *P* value. Collectively, these analyses demonstrate extensive transcriptional remodeling between established HFrEF at week 5 and the post-induction cannabidiol-treated state at week 9, involving metabolic, extracellular matrix, inflammatory, proliferative, and stress-response programs. Because the comparison includes differences in both treatment exposure and experimental time point, these transcriptional changes should be interpreted as reflecting the combined effects of recovery and post-induction cannabidiol treatment rather than a cannabidiol-specific effect alone. DEG indicates differentially expressed gene; GO, Gene Ontology; GSEA, gene set enrichment analysis; HFrEF, heart failure with reduced ejection fraction; MSigDB, Molecular Signatures Database; NES, normalized enrichment score; and pi-cannabidiol, post-induction synthetic cannabidiol treatment.

**Supplementary Figure 9. Concentration-dependent effects of synthetic cannabidiol on HUVEC morphology and metabolic activity. A**, Representative brightfield images of human umbilical vein endothelial cells (HUVECs) under untreated control, ethanol vehicle control (EtOH), and synthetic cannabidiol treatment at 0.1, 1, 2, 5, 10, and 20 μmol/L. HUVECs exposed to lower cannabidiol concentrations (0.1–2 μmol/L) maintained morphology broadly comparable to untreated and vehicle control conditions, whereas progressively higher concentrations were associated with reduced apparent cell density and morphological alterations. **B**, Quantification of cellular metabolic activity using the MTS assay, normalized to the untreated control condition. Synthetic cannabidiol produced a concentration-dependent reduction in MTS signal at higher concentrations, whereas lower concentrations exerted comparatively limited effects. Data were analyzed using the Kruskal–Wallis test followed by Dunn multiple-comparisons test because the data did not satisfy the normality assumption according to the Shapiro–Wilk test. *P*<0.05 was considered statistically significant. EtOH indicates ethanol; HUVEC, human umbilical vein endothelial cell; and MTS, 3-(4,5-dimethylthiazol-2-yl)-5-(3-carboxymethoxyphenyl)-2-(4-sulfophenyl)-2H-tetrazolium.

