## Supplementary Figures, Tables and Data fiels for "Synthetic Cannabidiol Attenuates Heart Failure Progression with Concomitant and Post Injury Administration Through Modulation of Immune and Endothelial to Mesenchymal Transition – Related Remodeling Programs": Supplemental Table1.pdf

**Supplementary Table 1:** Biological module/mechanisms linking the differentially expressed genes (DEGs) from the comparison between W 5HFrEF mice and the W5 HFrEF+c-s-cannabidiol groups in relation to literature available describing mechanisms of action of cannabidiol. The genes listed are all DEGs that were defined using **|log2FC| ≥ 1 and adjusted p < 0.05.** Genes were grouped into biological modules based on published cardiovascular and CBD-related literature, GO/MSigDB context, and biological plausibility. Because this is bulk whole-heart RNA-seq, the modules should be interpreted as hypothesis-generating whole-tissue signatures rather than cell-type-specific mechanisms.

| Biological module | Cannabidiol-upregulated DEGs | Cannabidiol-downregulated DEGs | Biological interpretation | References |
|---|---|---|---|---|
| **Activated fibroblast / ECM remodeling** | **Thbs2**, Spon2, Efemp1, Col6a6, Col20a1, Sned1, Adamts4, Ccbe1 | **Postn**, **Thbs4**, **Spp1**, Mfap4, Mfap5, Col8a1, **Scx**, Tbx15, Fstl3, Prrx2 | Cannabidiol downregulates several ECM / matricellular / fibroblast-remodeling genes, especially **Postn**, **Spp1**, **Scx**, **Thbs4**, **Mfap4**, and **Mfap5**, while upregulating **Thbs2**. This supports reduced pathological ECM remodeling rather than broad matrix shutdown. | **Postn:** ablation of periostin-expressing activated cardiac fibroblasts after chronic AngII exposure reduced fibrosis and improved cardiac function[1]. **Scx:** scleraxis governs fibroblast activation in pressure-overload HF; knockout attenuated fibrosis and improved function/survival[2]. **Thbs2:** TSP2 is required for myocardial matrix integrity under loading stress; TSP2-null mice exposed to AngII developed rupture/severe dysfunction with increased MMP activity[3]. **Spp1:** SPP1/osteopontin is linked to cardiovascular inflammatory and fibrotic remodeling, but its role is context-dependent[4]. |
| **Vascular / endothelial-associated guidance and remodeling — exploratory** | **Ccbe1**, Stab2, Slit2, **Slit3**, Sema3a, Adam19, Epha4, Ptpru, Celsr2 | **Apold1**, Nox4, Col8a1, Thbs4 | Cannabidiol alters genes with vascular, endothelial, angiogenesis/lymphangiogenesis, or guidance-related annotations. This should be described as **vascular-associated remodeling**, not direct evidence of endothelial rescue. Directionality is mixed and some genes are context dependent. | **Ccbe1:** CCBE1 is important for lymphatic endothelial budding, migration, and sprouting and enhances VEGF-C lymphangiogenic activity[5]. **Apold1:** APOLD1 is a vascular gene involved in pathological angiogenesis and endothelial homeostasis; cannabidiol downregulates it here, so interpretation should remain cautious[6]. **Slit3:** SLIT3 deficiency attenuated pressure-overload-induced fibrosis and hypertrophy, so cannabidiol-associated **Slit3 upregulation** is not straightforwardly |

| Biological module | Cannabidiol-upregulated DEGs | Cannabidiol-downregulated DEGs | Biological interpretation | References |
|---|---|---|---|---|
| | | | | protective and should not be overinterpreted[7]. |
| **Immune / inflammatory remodeling** | Cd1d1, Retnla, Ly75, Scara5, Clec18a, Ccl7, Tox, Zc3hav1l | Cd80, **Cxcr6**, Clec4n, **Aif1**, Trem2, Tmem119, Ms4a6b, Ms4a6c, Tlr13, Morrbid, Nlrc3, Serpinb1c | Cannabidiol changes immune-associated genes in both directions. The downregulated set includes several myeloid/macrophage or immune-activation-associated genes. This supports **immune-state remodeling**, not simple global immune suppression. | **Cxcr6:** CXCR6 deficiency attenuated pressure-overload-induced cardiac dysfunction/remodeling in mice, making cannabidiol-associated **Cxcr6 downregulation** directionally compatible with protection[8]. **Aif1:** AIF1 is linked to macrophage activation; macrophage AIF1 was reported to promote maladaptive inflammation and remodeling after ischemic cardiac injury, and knockdown improved repair[9]. **Trem2:** TREM2 macrophage biology in the heart is complex and can be reparative in MI, so cannabidiol-associated **Trem2 downregulation** should be interpreted as possible altered myeloid burden/state, not necessarily direct benefit[10]. |
| **Macrophage–fibrosis interface** | Retnla, Scara5, Ly75, Cd1d1, Ccl7 | Aif1, Cd80, Cxcr6, Clec4n, Trem2, Tmem119, Ms4a6b, Ms4a6c | The directional changes could suggest cannabidiol may alter macrophage/myeloid populations or activation states that interact with fibroblast remodeling. However, because several markers are cell-state dependent, this should remain a cautious bulk-RNA interpretation. | AngII-mediated cardiac fibrosis can require recruitment of myeloid fibroblast precursor cells into the heart, supporting a macrophage/fibroblast connection in AngII remodeling[11]. Cardiac macrophages can either promote repair or fibrosis depending on origin and activation state, so bulk RNA cannot define whether this module is uniformly protective[12]. |
| **Oxidative stress / cellular injury-response resolution** | Gsta3, Aldh1l2, Cox6b2, Hmgcs2, Acot1, Acot3 | **Nox4**, Hspa1a, Hspa1b, Atf3, Gdf15, Nupr1, Mt2, Car9, Ifi27l2a | Cannabidiol downregulates several stress-response genes and **Nox4**. This is consistent with reduced ongoing oxidative/injury signaling, but **Nox4** is context-dependent in | **Nox4:** cardiac-specific Nox4 deletion protected against pressure-overload injury in one study, but other work suggests endothelial Nox4 can protect against AngII-induced fibrosis; therefore, cannabidiol-associated **Nox4 downregulation** is plausible but not unambiguous[13]. |

| Biological module | Cannabidiol-upregulated DEGs | Cannabidiol-downregulated DEGs | Biological interpretation | References |
|---|---|---|---|---|
| | | | cardiovascular disease, so the claim is conservative. | Cannabidiol has published cardiac evidence for reducing oxidative/nitrative stress and remodeling in cardiomyopathy models[14]. |
| **Cardiomyocyte stress / hypertrophy / contractile remodeling** | Phkg1, Cox6b2 | **Acta1**, Ankrd2, **Fhl1**, Jsrp1, Synpo2l, Gdf15, Nmrk2, Hspa1a, Hspa1b | Cannabidiol downregulates several cardiomyocyte stress, cytoskeletal, or hypertrophy-associated genes, but individual genes are not specific to cardiomyocytes in the bulk RNA sequencing. | **Fhl1:** Fhl1 deficiency blunted pressure-overload hypertrophy and early HF progression while preserving function[15]. **Acta1:** skeletal α-actin is part of the fetal gene program in the heart; lower **Acta1** may be consistent with reduced fetal/stress remodeling[16]. cannabidiol protected AngII/L-NAME HF mice by reducing fibrosis/hypertrophy and preserving mitochondrial function, $Ca^{2+}$ handling, redox balance, and contractile function[14]. |
| **Membrane signaling / ion transport / excitable-cell biology** | Scn4b, Scn10a, Kcna1, Kcnv2, Kcnk1, Nalcn, Cnga3, Cngb3, Clcn1, Ano5, Aqp4, Chrna2, Grin2d, Adcy8, Adcy9 | Slc26a7, Aqp8, Kif1a, Tubb3, Ngef, Syndig1 | These genes while enriching in the neuronal mechanisms in the GO list, they likely reflect membrane excitability, ion handling, and cell-communication remodeling in the whole heart. | Cannabidiol's published AngII/L-NAME HF mechanism includes preserved cardiomyocyte $Ca^{2+}$ handling and excitation–contraction function, making this module biologically compatible with cannabidiol protection, although the individual genes listed require validation[14]. |
| **Cell migration / guidance / adhesion / tissue-patterning genes** | Epha4, Slit2, Slit3, Sema3a, Chl1, Reln, Gfra1, Gfra2, Nrxn2, Pcdhga4, Pcdhga12, Pcdhgb1, Pcdhgb5, Lrrc4b, Gpm6a, Colq | Ngef, Pcdhb9, Kif1a, Tubb3, Syndig1 | These genes probably contribute to axon-guidance / neuron-projection GO annotations. In cardiac bulk tissue, this should be described as guidance, adhesion, migration, or cell-communication biology rather than a primary nervous-system mechanism. | **Slit3** is the strongest cardiac-relevant gene in this category, but its direction is cautionary because Slit3 deficiency protected against pressure-overload fibrosis and hypertrophy in mice[7]. |

| Biological module | Cannabidiol-upregulated DEGs | Cannabidiol-downregulated DEGs | Biological interpretation | References |
|---|---|---|---|---|
| **Metabolic / lipid / mitochondrial remodeling** | Hmgcs2, Acot1, Acot3, Acsm5, Hsd11b1, Ces1d, Slc22a3, Aldh1l2, Alas2, Cox6b2 | Gck, Dio2, Star, Nmrk2, Car9 | Cannabidiol alters genes linked to lipid handling, ketone metabolism, steroid/thyroid-related pathways, mitochondrial enzymes, and hypoxia/metabolic stress. Directionality is mixed, so this should be framed as metabolic remodeling, not simple normalization. | Cannabidiol preserved mitochondrial membrane potential, respiratory control, oxidative balance, and bioenergetic function in the AngII/L-NAME HF model[14]. |

| Biological module | Cannabidiol-upregulated DEGs | Cannabidiol-downregulated DEGs | Biological interpretation | References |
|---|---|---|---|---|


7. Gong L, Wang S, Shen L, Liu C, Shenouda M, Li B, Liu X, Shaw JA, Wineman AL, Yang Y, Xiong D, Eichmann A, Evans SM, Weiss SJ and Si MS. SLIT3 deficiency attenuates pressure overload-induced cardiac fibrosis and remodeling. *JCI Insight*. 2020;5.
