## Supplementary Figures, Tables and Data fiels for "Synthetic Cannabidiol Attenuates Heart Failure Progression with Concomitant and Post Injury Administration Through Modulation of Immune and Endothelial to Mesenchymal Transition – Related Remodeling Programs": Supplementary Table 2.pdf

**Supplementary Table 2** Biological module/mechanisms linking the differentially expressed genes (DEGs) from the comparison between W5 HFrEF mice and the W5 HFrEF+c-s-cannabidiol groups in relation to literature available describing mechanisms of action of cannabidiol. A log2FC of at least 0.5 in either direction (up- or down-regulated) and a FDR adjusted p-value <0.05 was considered as significant for this analysis. Genes were grouped into predefined biological modules based on published cardiovascular biological plausibility. Because this is bulk whole-heart RNA-seq, the modules should be interpreted as hypothesis-generating whole-tissue signatures rather than cell-type-specific mechanisms.

| Predefined biological module | Curated genes altered by CBD (Gene, log2FC) | Biological interpretation | References |
|---|---|---|---|
| **EndMT-associated remodeling** | **Thbs4**↓ −1.46<br>**Bmp4**↓ −0.65<br>**Smad3**↑ +0.66<br>**Rbfox1**↑ +0.91<br>**Thbs2**↑ +1.17 | S-cannabidiol reduces a stress-remodeling / profibrotic endothelial–mesenchymal pressure signal through **Bmp4**↓ and **Thbs4**↓, while restoring genes linked to cardiac RNA-splicing adaptation and matrix integrity through **Rbfox1**↑ and **Thbs2**↑. **Smad3**↑ should be treated cautiously because total Smad3 mRNA does not prove SMAD3 pathway activation or suppression. | **Bmp4↓:** BMP4 impaired endothelial function and induced hypertension in mice partly by stimulating vascular NADPH oxidase activity, so its reduction is directionally compatible with reduced vascular oxidative stress.[1] **Thbs4↓:** THBS4 is induced in heart/vasculature under myocardial infarction, pressure overload, and hypertension; in AngII-treated Thbs4-null mice, hypertrophy, perivascular fibrosis/inflammation, and aortic dissections were reported, so lower Thbs4 with CBD is best interpreted as reduced upstream injury/remodeling pressure, not proof that THBS4 itself is harmful. [2] **Rbfox1↑:** RBFox1 is diminished in failing mouse and human hearts, and RBFox1 restoration attenuated pressure-overload hypertrophy, dysfunction, and fibrotic remodeling.[3] **Thbs2↑:** TSP2 is required for myocardial matrix integrity under loading stress; AngII-exposed TSP2-null mice developed cardiac rupture/dilatation with increased MMP activity.[4] **Smad3↑:** SMAD3 mediates fibrotic responses in pressure-overloaded heart, so total Smad3 mRNA increase is mechanistically ambiguous without phospho-SMAD3/localization data.[5] |

| Predefined biological module | Curated genes altered by CBD (Gene, log2FC) | Biological interpretation | References |
|---|---|---|---|
| **Endothelial phenotype** | **Kdr**↑ +0.71 | S-cannabidiol increases **Kdr/Vegfr2**, supporting preservation or restoration of an endothelial VEGFR2-associated program in the failing heart. This gene change is from bulk-heart RNA and should not be described as direct VEGFR2 activation (will need further validation with protein/phosphorylation or cell-localization data). | Cardiac endothelial VEGFR2 signaling induces angiogenesis and angiocrine release of ErbB receptor ligands that support endothelial–cardiomyocyte crosstalk and physiological cardiac growth.[6] |
| **Mesenchymal phenotype / fibroblast-state marker** | **Pdgfra**↑ +0.64 | S-cannabidiol increases **Pdgfra**, which is better interpreted here as restoration of a PDGFRα-associated resident fibroblast / stromal maintenance signal rather than as simple mesenchymal activation. This supports the reduced fibrosis seen in cannabidiol treated mice. | Loss of PDGFRα in cardiac fibroblasts caused rapid reduction of resident fibroblasts, supporting a role for PDGFRα signaling in cardiac fibroblast maintenance. [7] |
| **Fibrosis / matrix remodeling** | **Postn**↓ −1.49<br>**Serpine1**↓ −0.80<br>**Pdgfra**↑ +0.64<br>**Thbs2**↑ +1.17 | S-cannabidiol suppresses **Postn**, a marker of activated cardiac fibroblast remodeling, and reduces **Serpine1**, a fibrinolysis/ECM-regulatory gene, while increasing **Pdgfra** and **Thbs2**, consistent with a shift away from activated fibrotic remodeling toward fibroblast/matrix homeostasis. **Serpine1/PAI-1** should be interpreted cautiously because cardiac PAI-1 biology is context-dependent. | **Postn↓:** ablation of periostin-expressing activated cardiac fibroblasts after chronic AngII exposure reduced cardiac fibrosis and improved cardiac function.[8] **Thbs2↑:** TSP2 supports myocardial matrix integrity under loading stress. [4] **Pdgfra↑:** PDGFRα signaling supports resident cardiac fibroblast maintenance.[7] **Serpine1↓:** SERPINE1/PAI-1 regulates fibrinolysis and ECM-related protease activity, but in AngII/aldosterone hypertension models PAI-1 deficiency worsened cardiac fibrosis, so CBD-associated Serpine1 reduction should be framed as reduced remodeling/stress-associated expression rather than unequivocally antifibrotic by itself.[9] |

| Predefined biological module | Curated genes altered by CBD (Gene, log2FC) | Biological interpretation | References |
|---|---|---|---|
| **Inflammation / myeloid-associated remodeling** | **Lgals3**↓−0.79<br>**Lyz2**↓ −0.50 | S-cannabidiol reduces **Lgals3** and **Lyz2**, supporting reduced galectin-3/myeloid-associated inflammatory remodeling in the heart. This is compatible with reduced fibrosis, but bulk RNA cannot distinguish fewer myeloid cells from altered myeloid activation state. | **Lgals3↓:** genetic or pharmacologic inhibition of galectin-3 attenuated cardiac fibrosis, LV dysfunction, and heart-failure development in remodeling models.[10] **Lyz2↓:** Lyz2 encodes lysozyme 2 and is expressed in hematopoietic/myeloid-associated compartments; its decrease supports reduced myeloid-associated transcript burden but is not by itself a cardiac-fibrosis mechanism.[11] |
| **Inflammasome / inflammatory danger signaling** | **Panx1**↓ −0.96 | S-cannabidiol suppresses **Panx1**, supporting reduced pannexin-1-associated ATP/danger signaling. This is a plausible bridge between membrane-channel biology, inflammatory activation, and fibroblast activation. It does not prove canonical NLRP3 inflammasome inhibition (further studies of caspase-1, IL-1β maturation, GSDMD cleavage, ASC specks, or NLRP3 protein data are needed). | Cardiomyocyte ATP release through PANX1 channels was proposed to activate fibroblasts and drive early fibrosis after cardiac stress.[12] |
