## Supplementary figures and images for "Synthetic Cannabidiol Attenuates Heart Failure Progression with Concomitant and Post Injury Administration Through Modulation of Immune and Endothelial to Mesenchymal Transition – Related Remodeling Programs"

### Supplementary Figure 1.tif

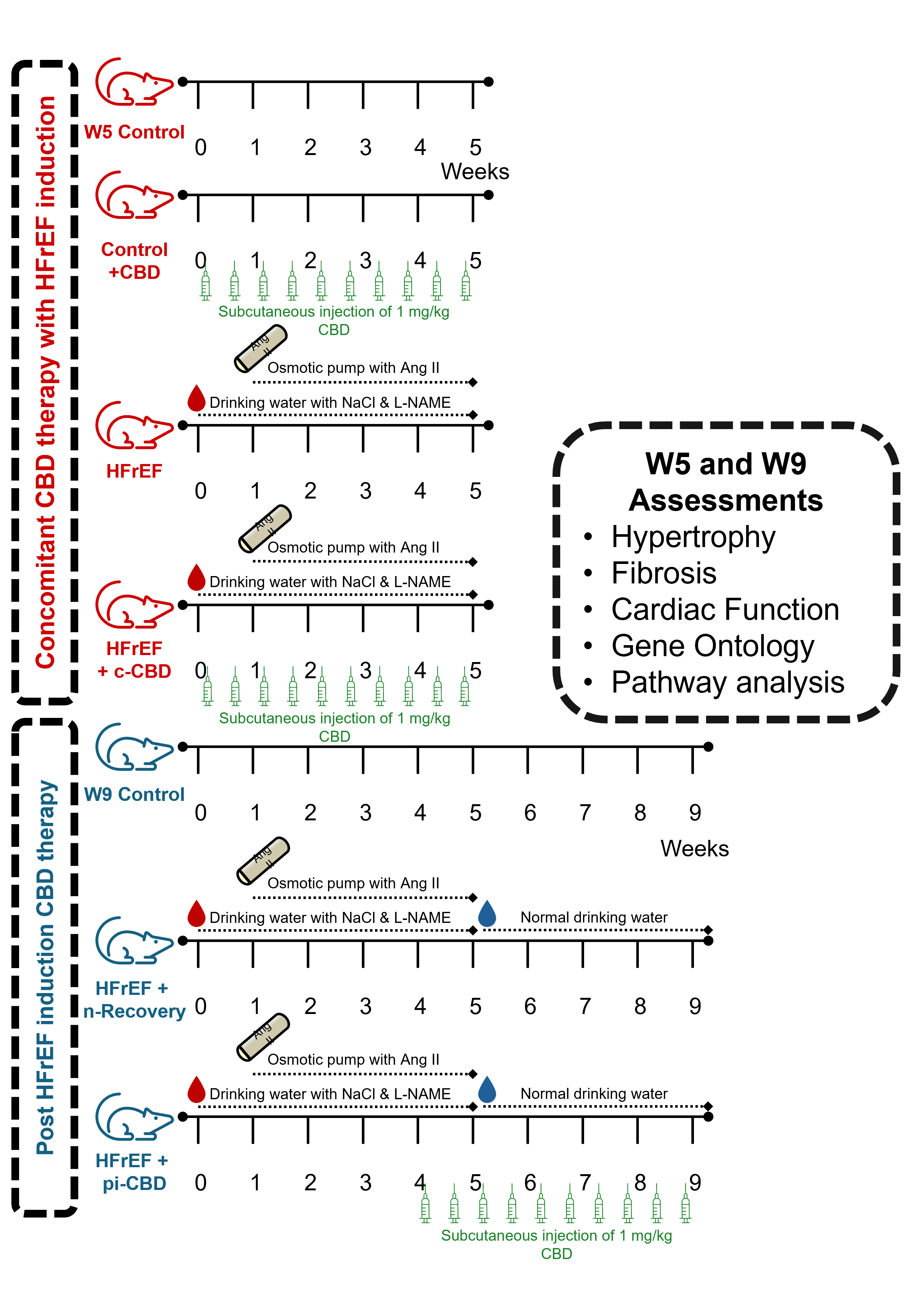

### Supplementary Figure 2.tif

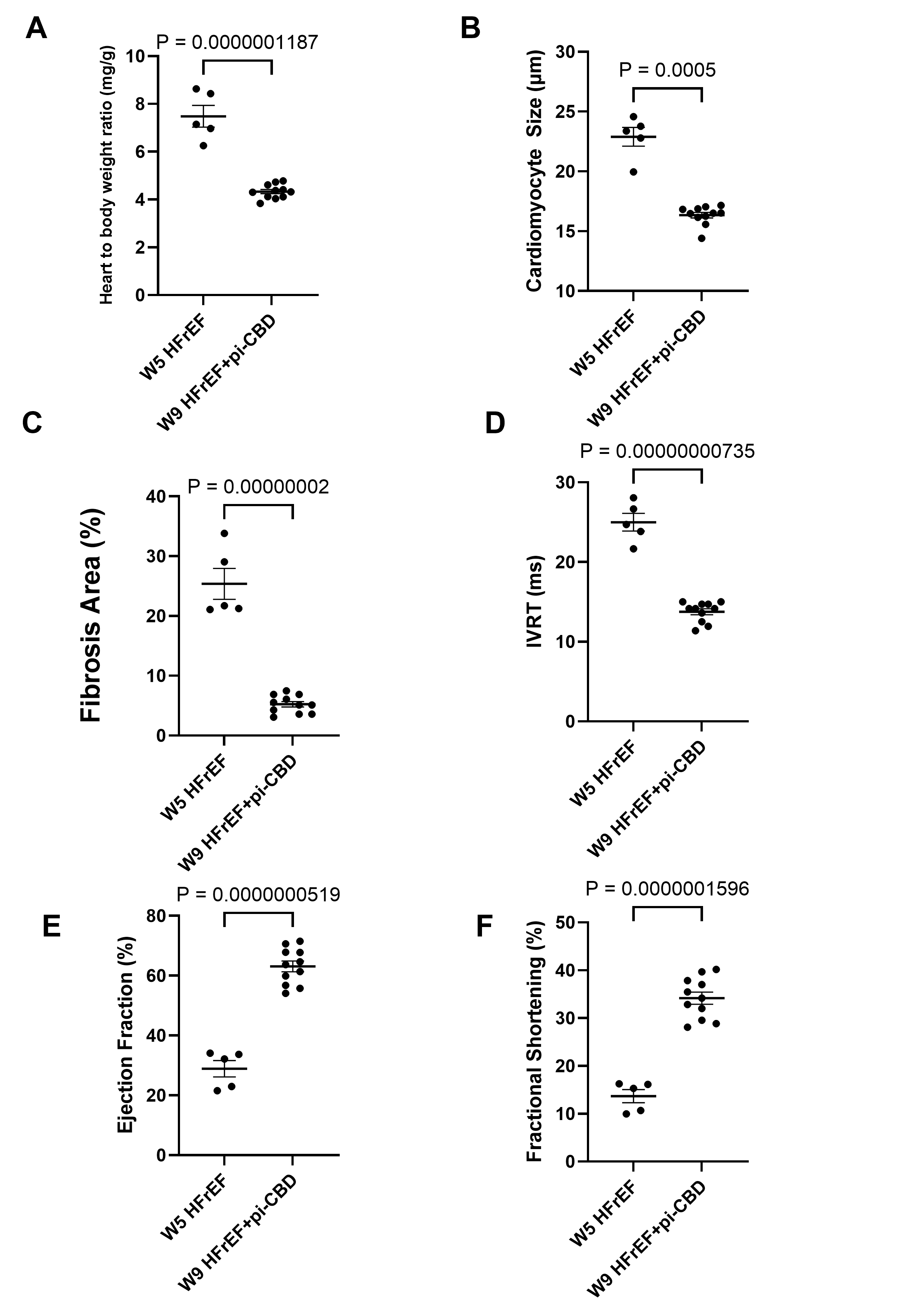

### Supplementary Figure 3.tif

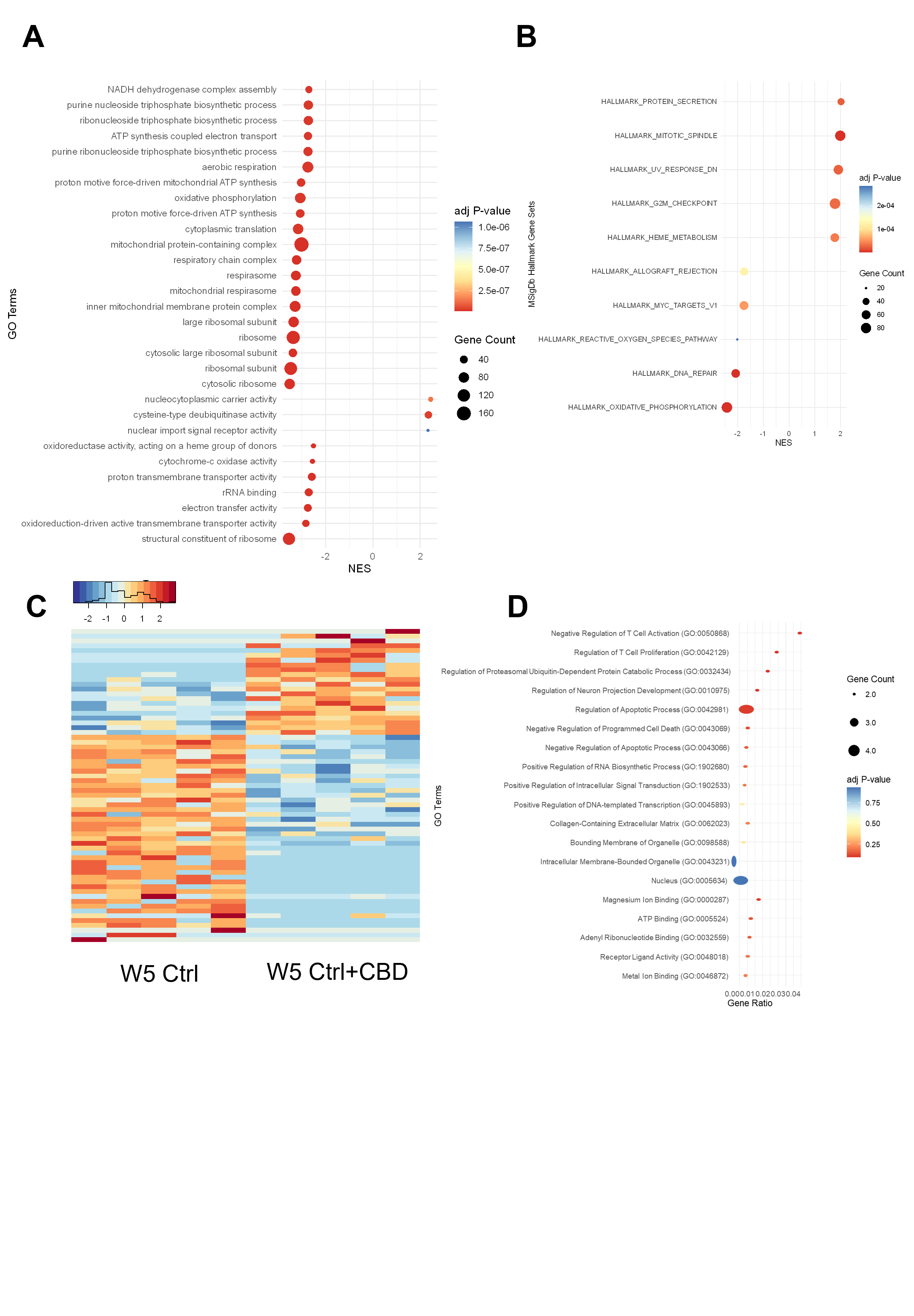

### Supplementary Figure 4.tif

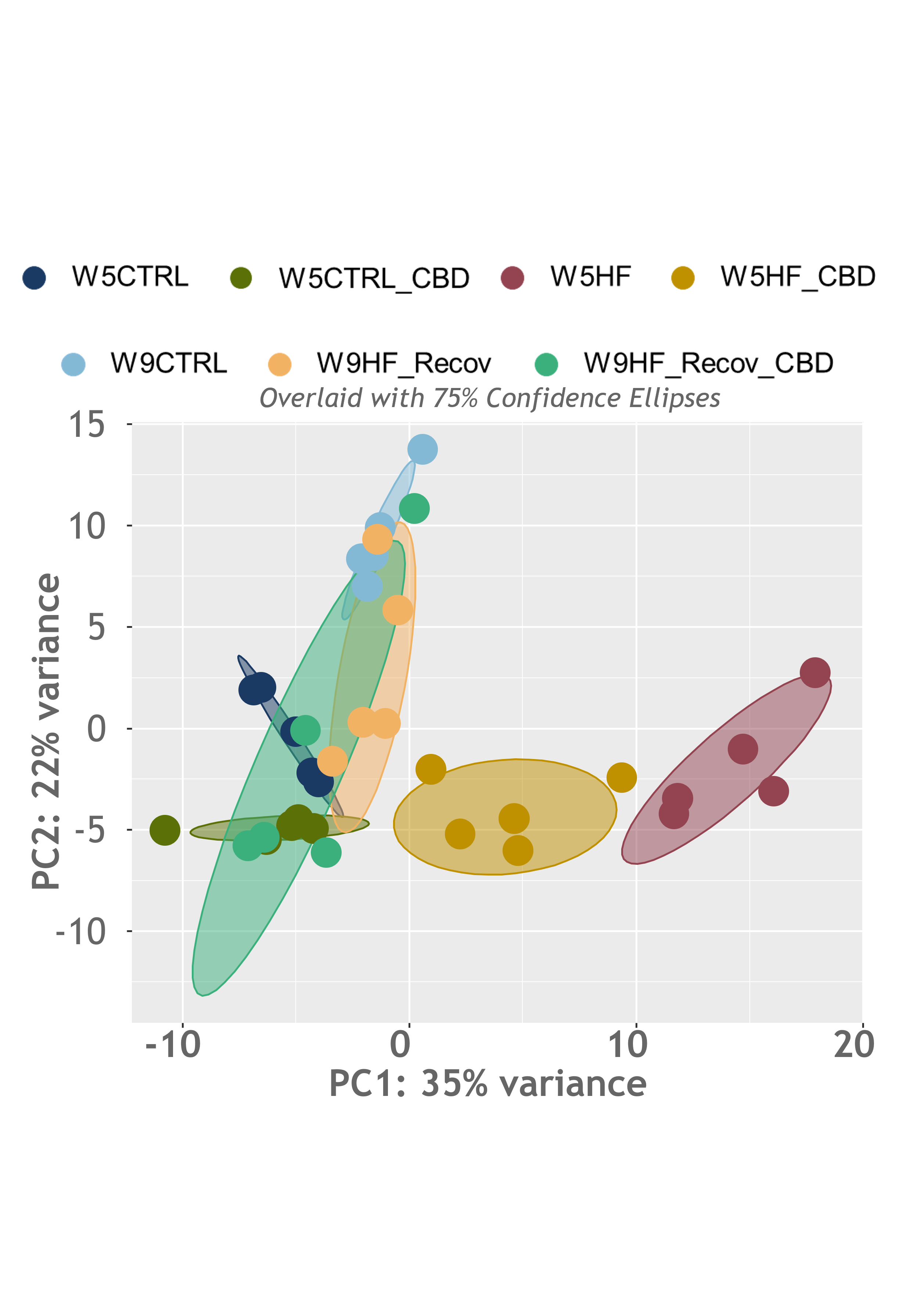

### Supplementary Figure 5.tif

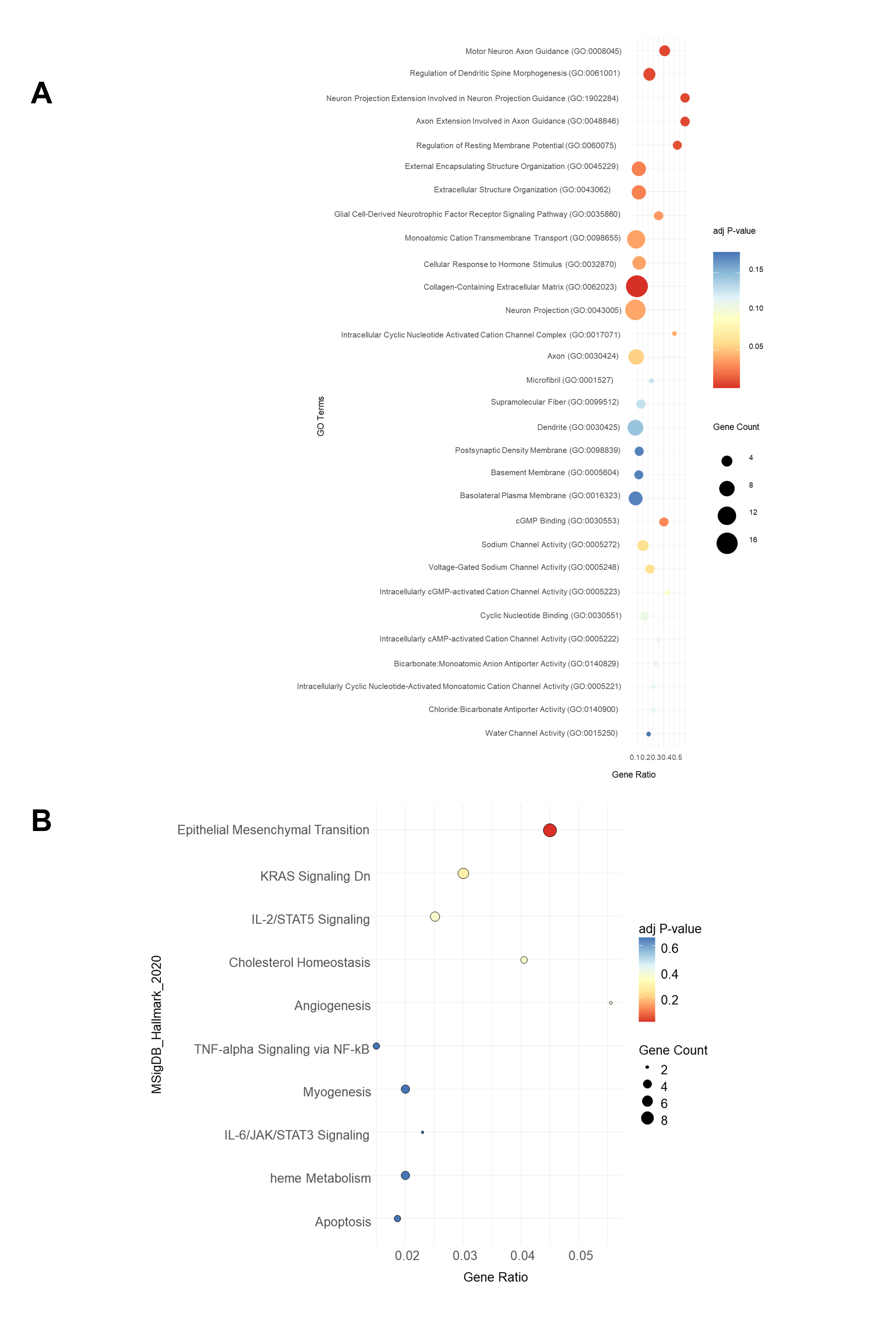

### Supplementary Figure 6.tif

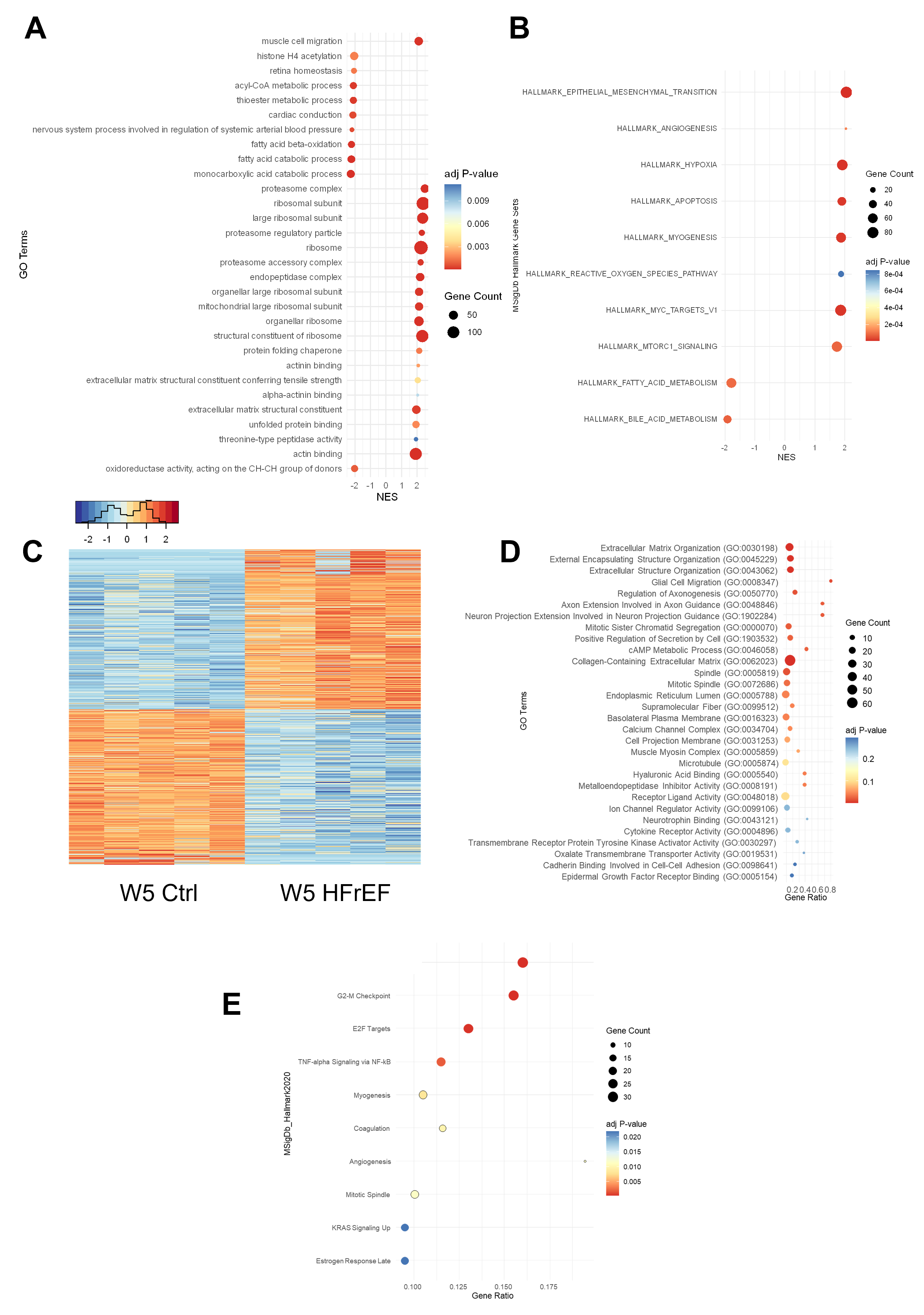

### Supplementary Figure 7.tif

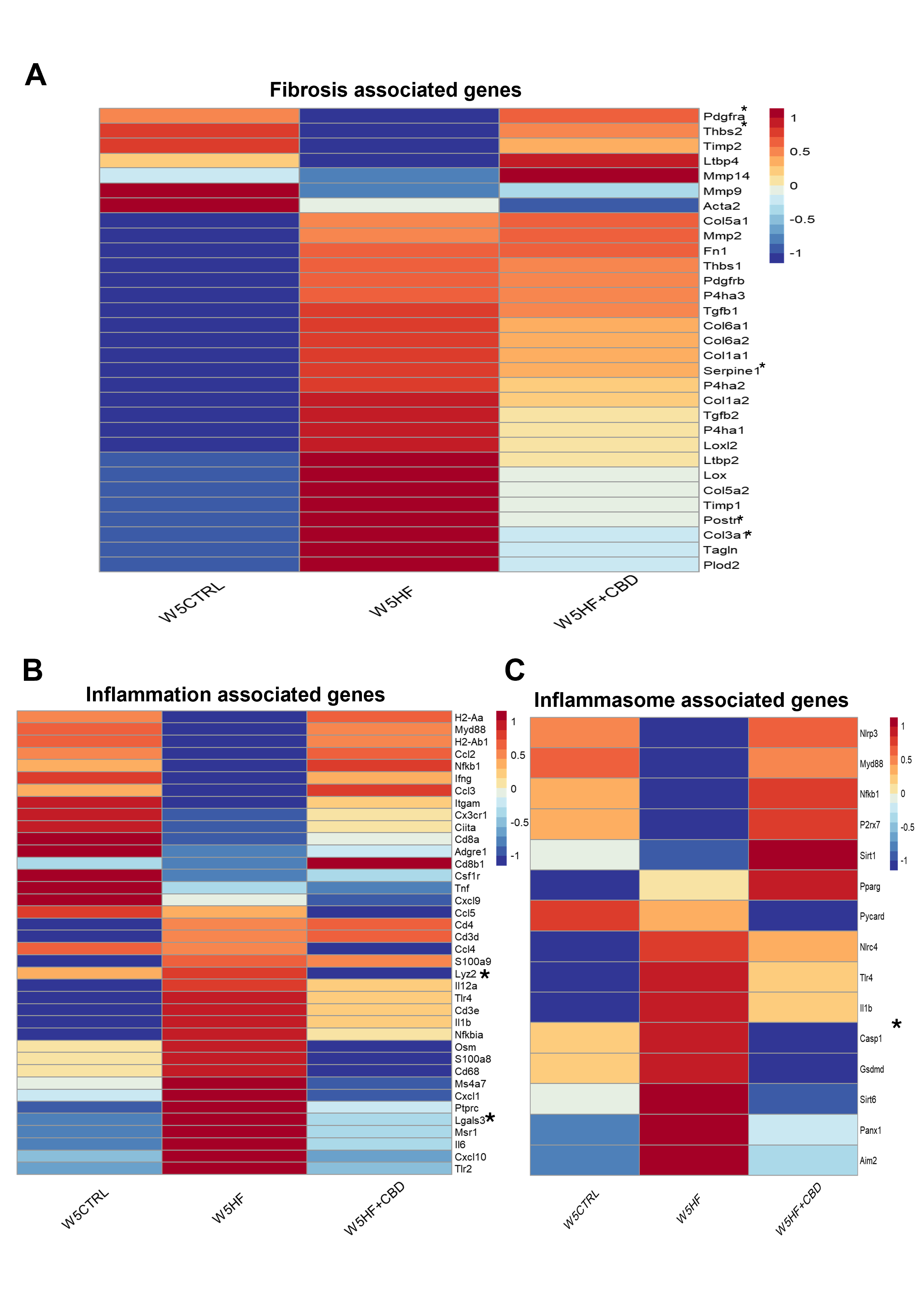

### Supplementary Figure 8.tif

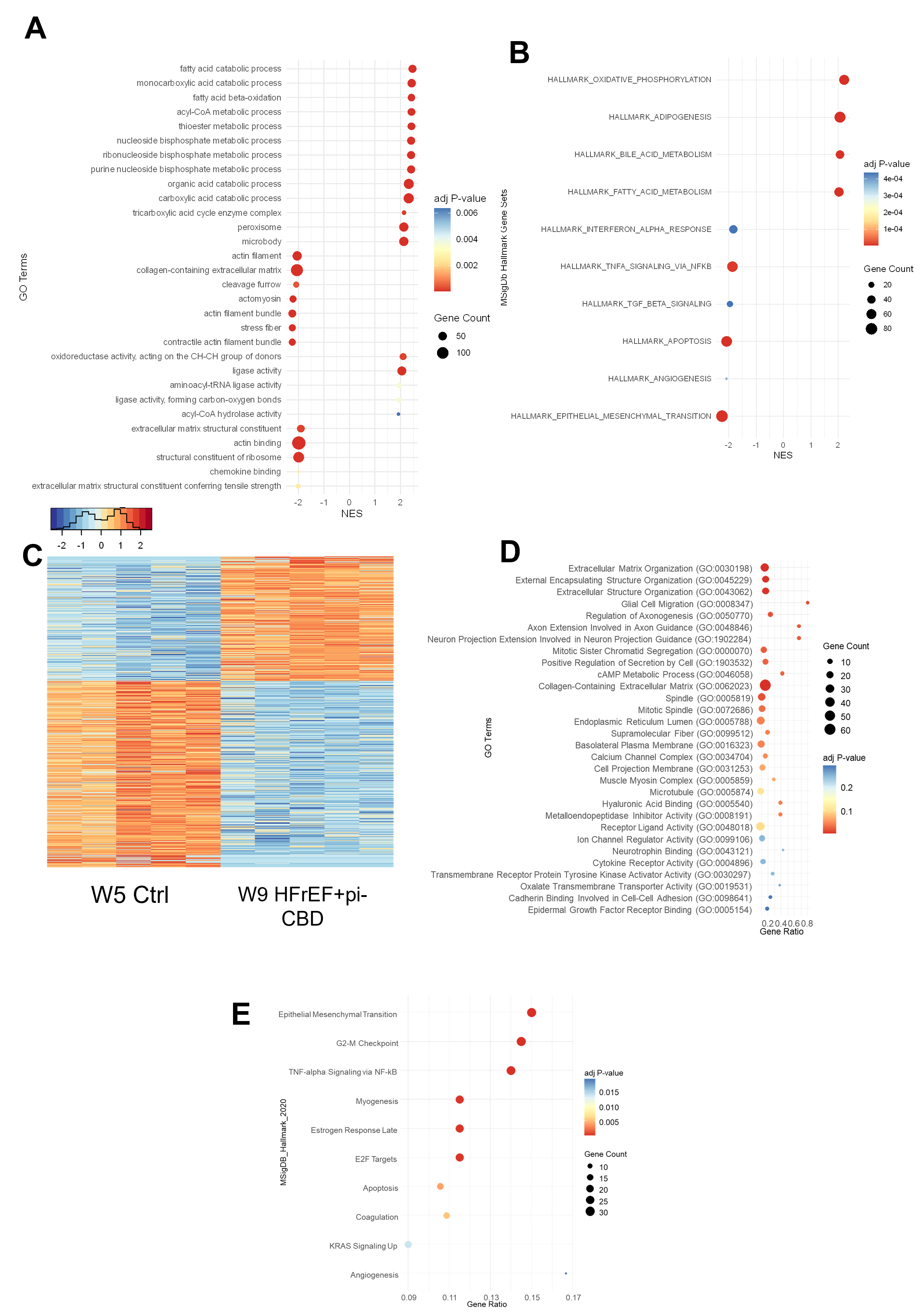

### Supplementary Figure 9.tif

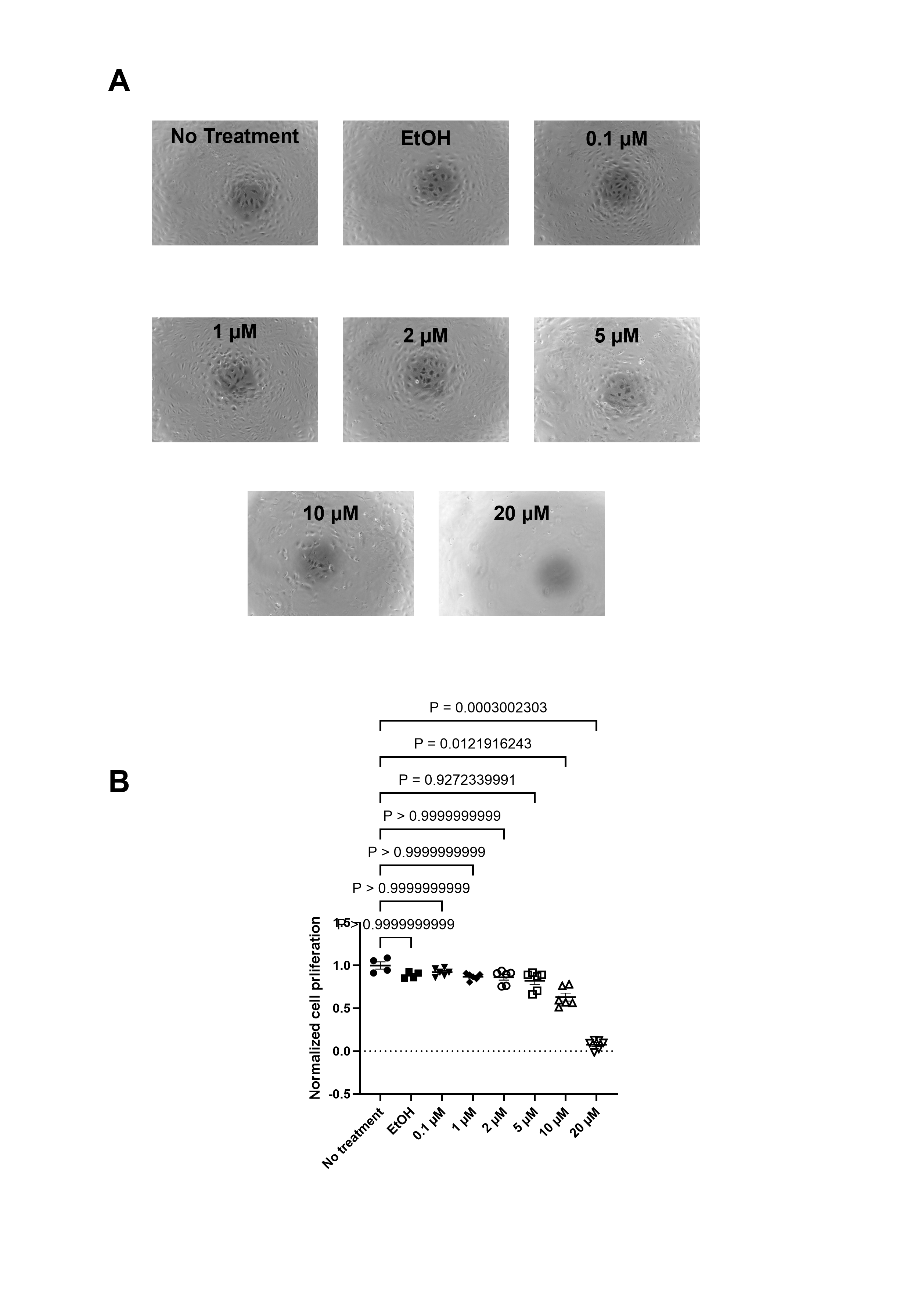
